# The combination of BCL-xL PROTAC and mTOR inhibitor sensitizes pancreatic ductal adenocarcinoma to KRAS^G12D^ inhibitor treatment by enhancing apoptosis induction

**DOI:** 10.64898/2026.01.05.697773

**Authors:** Javed Miyan, Vignesh Vudatha, Lin Cao, Peiyi Zhang, Guangrong Zheng, Lei Zheng, Jose Trevino, Daohong Zhou, Sajid Khan

## Abstract

Pancreatic ductal adenocarcinoma (PDAC) is a highly aggressive cancer with a five-year survival rate of approximately 13%, partly because of limited treatment options and resistance to therapies. Although the recently discovered KRAS G12D inhibitor MRTX1133 has shown promise for PDAC treatment in preclinical studies, its clinical efficacy as a single agent is expected to be limited, as is the case with KRAS G12C inhibitors. Therefore, in this study, we evaluated potential combination strategies to enhance the therapeutic effect of MRTX1133. We rationally combined MRTX1133 with the BCL-xL proteolysis-targeting chimera (PROTAC) DT2216 and the mTOR inhibitor everolimus, which significantly potentiated the anti-tumor activity of MRTX1133 in multiple G12D-mutated PDAC cells in vitro by enhancing apoptosis induction. Mechanistically, MRTX1133 treatment increased BIM and decreased NOXA levels, while the combination of DT2216/everolimus was shown to simultaneously enhance BIM release and stabilize NOXA. In vivo, DT2216/everolimus combination significantly potentiated the anti-tumor activity of MRTX1133 in AsPC1 PDAC xenograft model. Furthermore, the triple combination of MRTX1133, DT2216, and everolimus effectively overcame acquired resistance to MRTX1133 in AsPC1 cells in vitro and in xenograft model. Collectively, our findings suggest that the single-agent efficacy of MRTX1133 is limited by apoptosis inhibition, and that its combination with DT2216/everolimus can enhance apoptosis and potentially overcome resistance in KRAS G12D-mutated PDAC.

**SIGNIFICANCE:** KRAS inhibitors, including KRAS G12D inhibitor MRTX1133, are promising therapeutics against KRAS-mutated pancreatic ductal adenocarcinoma (PDAC), but drug resistance limits their efficacy. Our study reveals that robust induction of apoptosis using a combination of BCL-xL PROTAC degrader and an mTOR inhibitor, significantly enhances MRTX1133 efficacy in PDAC models without increasing toxicity to normal tissues.

## INTRODUCTION

Pancreatic ductal adenocarcinoma (PDAC) comprises >90% of pancreatic cancer cases, is highly aggressive, and has one of worst prognoses among human cancers.^1^ The recent discoveries of KRAS inhibitors (KRASi), including pan-KRAS inhibitors and allele-specific KRAS inhibitors (including KRAS G12C inhibitors (G12Ci) and KRAS G12D inhibitors (G12Di)) have sparked renewed optimism for treating PDAC patients whose tumors harbor KRAS mutations. Among these inhibitors, G12Di MRTX1133 could particularly be a promising treatment for PDAC patients.^2–6^ This is because G12D mutations occur in ∼45% of PDAC patients, and are associated with poorer prognosis compared to non-KRAS oncogenic drivers.^7^ Though MRTX1133 is yet to be evaluated in clinical trials, its clinical efficacy as a monotherapy in PDAC is likely to be limited, as observed with G12Ci, due to myriad of intrinsic, adaptive and acquired resistance mechanisms.^8–11^ Among these mechanisms, epithelial-to-mesenchymal transition (EMT) has been previously reported as a key contributor to KRASi resistance, particularly in treatment refractory PDAC models.^12–14^ In addition, the demonstrated mechanisms of resistance to G12Ci, such as feedback and compensatory activation of receptor tyrosine kinase (RTK)-mediated mitogen-activated protein kinase (MAPK) and the PI3K-AKT-mTOR signaling pathway, have also been shown to be involved in resistance to MRTX1133 in early preclinical studies.^4, 15–17^ However, targeting effectors of these pathways individually is insufficient to overcome KRASi resistance because multiple RTKs are involved in the reactivation of various downstream effectors. This results in heterogenous and short-lived responses by targeting a single RTK or MAPK effector in combination with KRASi, in part because of redundancy in KRAS downstream signaling effectors.^15–17^ Though targeting more than one of the effectors of these pathways in combination with MRTX1133 might improve efficacy, which may also lead to higher normal tissue toxicities limiting the potential for clinical translation. Therefore, more effective and safer combination strategies to enhance the initial efficacy of and overcome resistance to MRTX1133 are necessary.

We and others have recently shown that the inability of G12Ci monotherapy to induce robust cell death, due to overexpression of BCL-xL, is one of the underlying mechanisms of resistance.^18, 19^ BCL-xL is an important anti-apoptotic cancer target protein that belongs to BCL-2 family and is significantly upregulated in human PDACs compared to normal pancreas tissue.^20^ Its expression is negatively correlated with PDAC patient survival and response to therapy.^20^ However, targeting BCL-xL with conventional inhibitors causes thrombocytopenia, an on-target and dose-limiting toxicity.^21–23^ To overcome thrombocytopenia, we have developed DT2216, a platelet-sparing BCL-xL proteolysis-targeting chimera (PROTAC) degrader, by linking navitoclax (a BCL-xL/BCL-2 dual inhibitor) with a ligand for Von Hippel-Lindau (VHL) E3 ligase. DT2216 targets BCL-xL to the VHL for ubiquitination and proteasomal degradation of the former.^24^ Because VHL is minimally expressed in platelets but abundantly expressed in most tumor cells, DT2216 is highly potent against tumor cells when used alone or in combination with other antitumor agents and has limited antiplatelet activity compared to navitoclax.^18, 24–28^ Based on these preclinical findings, DT2216 has received orphan drug designation and fast-track designation from the FDA. DT2216 has completed a phase I clinical trial in patients with relapsed or refractory solid malignancies (NCT04886622), where it was generally well tolerated. Most patients experienced only transient thrombocytopenia with rapid recovery and demonstrated BCL-xL depletion in peripheral leukocytes.^29^ Based on findings from this trial and promising preclinical results, DT2216 has advanced to a phase I/II clinical trial in combination with irinotecan in relapsed/refractory fibrolamellar carcinoma (NCT06620302), and phase I trial in combination with paclitaxel in platinum resistant ovarian cancer (NCT06964009).

In this study, we hypothesize that endogenous and MRTX1133-induced dysregulation of BCL-2 family proteins (including BCL-xL) coupled with mTOR activation can contribute to intrinsic and acquired MRTX1133 resistance, respectively, through evasion of apoptosis. Therefore, we aimed to develop a rational combination strategy to enhance the efficacy of G12Di in PDAC. Specifically, we used DT2216 and everolimus (an FDA-approved mTOR inhibitor (mTOR-i)) to overcome intrinsic and acquired resistance to MRTX1133 by augmenting tumor cell apoptosis in G12D-mutated PDAC cell lines and xenograft models. Our findings suggest that MRTX1133 induces apoptotic reprogramming by induction of pro-apoptotic BIM and suppression of pro-apoptotic NOXA. Furthermore, these reprogrammed cells are vulnerable to DT2216 and/or everolimus for induction of apoptosis via degrading BCL-xL, thereby releasing BIM, and upregulation of NOXA, respectively. These agents work synergistically to improve antitumor efficacy of MRTX1133 not only *de novo* but also in G12Di resistant PDAC *in vitro* and *in vivo*.

## MATERIAL AND METHODS

### Cell lines and cell culture

Human PDAC cell lines (AsPC1 (RRID:CVCL_0152), HPAC (RRID:CVCL_3517), SW1990 (RRID:CVCL_1723), PANC1 (RRID:CVCL_0480) and BxPC3 (RRID:CVCL_0186)) were purchased from the American Type Culture Collection (ATCC, Manassas, VA). AsPC1, SW1990 and BxPC3 cells were cultured in RPMI-1640 medium (Cat #22400–089, Thermo Fisher, Waltham, MA). PANC1 cells were cultured in Dulbecco’s modified Eagle’s medium (DMEM) (Cat #12430-062, Thermo Fisher). All culture media were supplemented with 10% heat-inactivated fetal bovine serum (FBS) (Cat #S11150H, Atlanta Biologicals, GA), 1% penicillin-streptomycin (Pen-Strep) solution (Cat #15140122, Thermo Fisher). HPAC cells were cultured in DMEM/F-12 medium (Cat #11330032, Thermo Fisher) supplemented with 5% FBS, 0.002 mg/ml insulin (Cat #12585-014, Thermo Fisher), 0.005 mg/ml human transferrin (Cat #T8158, MilliporeSigma, Burlington, MA), 40 ng/ml hydrocortisone (Cat #H6909, MilliporeSigma), 10 ng/ml mouse epidermal growth factor (Cat #CB-40010, Fisher Scientific) and 1% pen-strep. Cell lines were used for experiments within 10 passages after thawing from liquid nitrogen. All cultures were confirmed for Mycoplasma negativity using the MycoAlert Mycoplasma Detection Kit (Cat #LT07–318, Lonza, Basel, Switzerland). All the cell lines were maintained in a humidified incubator at 37° C and 5% CO_2_.

### Chemical compounds

For *in vitro* experiments, DT2216 was provided by Dr. Guangrong Zheng’s laboratory (University of Florida, Gainesville, FL), which they synthesized according to the previously described protocol.^24^ For *in vivo* experiments, research grade DT2216 was kindly provided by Dialectic Therapeutics (Dallas, TX). MRTX1133 (Cat #HY-134813) and everolimus (Cat #HY-10218) were purchased from MedChemExpress (Monmouth Junction, NJ). For *in vitro* experiments, the compounds were dissolved in DMSO at 10 mM stock solution. *In vivo* formulations are described in the xenograft study section.

### Cell viability assays

Cells were seeded in CellStar µClear white 96-well plates (Cat #5665-5098, USA Scientific, Ocala, FL) at a density of 5,000 cells per well. After overnight incubation, different concentrations of the drugs were added in triplicates to the plates. The PrestoBlue™ HS Cell Viability Reagent (Cat #P50201, Thermo Fisher) was used to measure cell viability according to the manufacturer’s protocol. The fluorescence intensity was recorded at 560 nm (excitation)/590 nm (emission) using Synergy Neo2 multimode plate reader (Biotek, Winooski, VT).

### Colony formation assays

Cells were seeded in 6-well plates at a density of 2,000 cells per well. After overnight incubation, the cells were treated with test compounds for 10-14 days. At the end of treatment, the cells were fixed with absolute methanol, and then stained with 0.1% crystal violet solution. The images were captured using ChemiDoc MP Imaging System (Bio-Rad, Hercules, CA). Thereafter, 1 mL of 10% acetic acid solution was added per well to dissolve stained-colonies, and the absorbance was recorded at 590 nm as a direct measure of colony growth.

### Annexin V/PI staining and flow cytometric analysis

Cells were harvested by trypsinization, including dead cells from the culture medium, and centrifuged at 300 × g for 5 min at 4 °C. The cell pellet was washed twice with cold PBS and resuspended in 1× Annexin V Binding Buffer (Cat #422201, BioLegend, San Diego, CA) at a concentration of 2.5 × 10^6^ cells/mL Approximately, 2.5 × 10^5^ cells in 100 µL suspension were transferred into flow cytometry tubes. Cells were stained with Alexa Fluor 647-conjugated AnnexinV (Cat # 640911, BioLegend) and propidium iodide (PI). Samples were incubated for 30 min at room temperature in the dark, followed by dilution with 400 µL Annexin V Binding Buffer. Data acquisition was performed on a BD FACSCelesta flow cytometer. Apoptotic populations (live, early apoptotic, late apoptotic/necrotic) were analyzed based on Annexin V and PI staining using FlowJo software (Tree Star Inc.).

### Immunoblotting

Cells were lysed in RIPA buffer (Cat #BP-115DG, Boston Bio Products, Ashland, MA) supplemented with protease + phosphatase inhibitor cocktail (Cat #PPC1010, MilliporeSigma) and the assay was performed as described previously.^24^ Briefly, an equal amount of proteins (∼40 µg/lane) were loaded to a precast gel and transferred onto PVDF membranes. The membranes were blocked with 5% (w/v) non-fat dry milk in TBS-T buffer, and subsequently probed with primary antibodies overnight at 4 °C. After washing with TBST, the membranes were incubated with horseradish peroxidase (HRP)-linked secondary antibody for 1 h at room temperature. Finally, the membranes were incubated with chemiluminescent HRP substrate (Cat #WBKLS0500, MilliporeSigma), and imaged using the ChemiDoc MP Imaging System. The densitometric analysis of immunoblots was performed using Image J software (RRID:SCR_003070). The details of antibodies are provided in Supplementary Table-1.

### RNA extraction and quantitative real-time PCR

RNA was isolated from cells using RNeasy Mini Kit (Cat #74106, Qiagen, Hilden, Germany). A total of 1 µg of RNA was converted into cDNA using high-capacity cDNA reverse transcription kit (Cat #4368813, Applied Biosystems, Foster City, CA) as per the manufacturer’s instructions. The mRNA expressions were quantified using pre-designed Taqman probes of *BCL2L11* (Assay #Hs00708019_s1) and *PMAIP1* (Assay # Hs00560402_m1) from Thermo Fisher. The expression of *GAPDH* (Assay #Hs02786624_g1) was used for normalization and the level of gene expression in untreated cells was used as a baseline. Fold change in gene expression was calculated using the ΔΔCT method.

### Immunoprecipitation

Cells were lysed in the Pierce IP lysis buffer (Cat #87787; Thermo Fisher) supplemented with protease + phosphatase inhibitor cocktail and the assay was performed as described previously.^24^ The supernatants were collected and precleared by incubating with 1 µg of mouse anti-IgG (Cat #sc-2025; Santa Cruz Biotechnology, Dallas, TX) and 20 µL of protein A/G-PLUS agarose beads (Cat #sc-2003; Santa Cruz Biotechnology) for 30 min at 4 °C. The supernatants containing 1 mg of protein were incubated with 2 µg of anti-BIM (Cat #sc-374358, Santa Cruz Biotechnology, RRID:AB_10987853), or anti-MCL-1 (Cat #sc-12756, Santa Cruz Biotechnology, RRID:AB_627915) or anti-IgG antibody (Cat # sc-2025, Santa Cruz Biotechnology, RRID:AB_737182) overnight followed by incubation with 25 µL protein A/G-PLUS agarose beads for 1 h at 4 °C. Thereafter, the protein A/G-PLUS agarose beads were collected by centrifugation, washed three times with IP lysis buffer, mixed with 50 µL of Laemmli’s SDS-buffer and boiled to release protein into the supernatant, which was then subjected to immunoblot analysis for BIM, BCL-xL, MCL-1, BAX, BAK and NOXA. Total cell lysate (input) samples were run alongside. Anti-rabbit HRP-conjugated Fc fragment specific secondary antibody (Cat #111-035-046, RRID:AB_2337939, dilution 1:10000, Jackson ImmunoResearch, West Grove, PA) was used to detect immune complexes in immunoblotting.

### Generation of MRTX1133 resistant cells

AsPC1 cells were initially exposed to 100 nM of MRTX1133, and after they became resistant, the concentration was increased to 200 nM. The concentration of MRTX1133 was kept on doubling until the cells became resistant to 3.2 µM of MRTX1133. Totally, it took two months for the cells to become resistant to 3.2 µM of MRTX1133. At this time, the cells were considered MRTX1133 resistant (referred to as AsPC1-MR) to be used for experiments and maintained in 3.2 µM of MRTX1133.

### Xenograft study

SCID-Beige female mice aged 5-6 weeks were purchased from the Charles River Laboratories (Stock No. 250, Wilmington, MA). AsPC1 cells mixed with 50% Matrigel (Cat #356237, Corning, Corning, NY) and 50% plain RPMI-1640 medium (5×10^6^ cells/100 µL/mouse) were injected subcutaneously (s.c.) into the right flank region as described previously.^24^ Tumor growth was monitored daily, and tumor size was measured twice a week with digital calipers and tumor volume was calculated using the formula (Length×Width^2^×0.5). The mice were randomized into different treatment groups when their tumors reached 150 mm^3^ in volume. For efficacy study, mice were treated with vehicle, MRTX1133 (3 mg/kg, bid, 5 days ON/2 days OFF, i.p.), DT2216 (15 mg/kg, twice a week, i.p.), everolimus (2.5 mg/kg twice a week, p.o.) MRTX1133 + DT2216, MRTX1133 + Everolimus and MRTX1133 + DT2216 + Everolimus. MRTX1133 and Everolimus were dissolved into one-part DMSO, followed by the addition of nine parts of 20% (v/v) Captisol (Cat. No. NC0604701, Cydex Pharmaceuticals, a Ligand Company, San Diego, CA, USA) pre-prepared in normal saline. DT2216 was formulated by a CRO. All the animal procedures were performed in accordance with the IACUC guidelines at the University of Texas Health Science Center at San Antonio.

MRTX1133-resistant xenograft model: The tumors were considered resistant when they showed significant regrowth from their maximum regression after initial MRTX1133 treatment at 10 mg/kg, bid, 7 days a week, ip. Consistent tumor growth despite continued MRTX1133 exposure confirmed the establishment of acquired resistance in this model. At this point, mice were randomized into four different treatment groups i.e., MRTX1133 (10 mg/kg, bid, 7 days a week, i.p.), MRTX1133 (10 mg/kg, bid, 7 days a week, i.p.) plus DT2216 (15 mg/kg, twice a week, i.p.), MRTX1133 (10 mg/kg, bid, 7 days a week, i.p.) plus everolimus (2.5 mg/kg twice a week, p.o.), and MRTX1133 (10 mg/kg, bid, 7 days a week, i.p.) plus DT2216 (15 mg/kg, twice a week, i.p.) plus everolimus (2.5 mg/kg twice a week, p.o.).

### Pharmacodynamic (PD) study

Tumor-bearing mice were treated twice daily (bid) for 5 days with MRTX1133, and two doses of DT2216 and everolimus at 3-day interval and same dosage as used in the xenograft study. For PD study, the treatment was started when average tumor size reached ∼400 mm^3^. The tumors were subsequently harvested 24 hours after the last dose of DT2216. A portion of tumors were flash frozen to be used for immunoblotting analysis, and another portion was fixed in 4% paraformaldehyde for histological analyses.

### Hematoxylin & Eosin (H&E) staining

H&E staining was performed using the Leica ST5010 Autostainer XL (Leica Biosystems, Deer Park, IL). Tissue sections were deparaffinized by incubating at 60 °C for 30 min, followed by four washes in xylene (5 min each). Sections were then rehydrated through a graded concentrations of ethanol (100%, 95%, 80%, and 70%, 5 min each) and rinsed in distilled water (dH_2_O). For nuclear staining, sections were incubated in hematoxylin, rinsed in dH_2_O, differentiated in an acidic alcohol solution, blued in an alkaline solution, and rinsed again in dH_2_O. Eosin was then applied for cytoplasmic staining. After staining, sections were dehydrated through graded ethanol (70% to 100%) and cleared in xylene (4x, 5 min each). Finally, slides were mounted using synthetic permount and a glass coverslip. Images were acquired using Zeiss Axio Vert.A1 inverted microscope under 10x magnification.

### Immunohistochemistry (IHC)

IHC staining was performed using the Leica ST5010 Autostainer XL and the Dako EnVision FLEX system (Agilent, Santa Clara, CA). Tissue sections were deparaffinized by heating at 60°C for 30 min, followed by xylene washes (4x, 5 min each) and rehydration through a graded ethanol series (100% ethanol twice for 5 min each, 95% ethanol for 5 min, and 70% ethanol for 5 min). Heat-induced epitope retrieval was carried out using Dako PT Link with EnVision FLEX Target Retrieval Solution (low or high pH) at 97 °C for 20 min. After retrieval, sections were washed in EnVision FLEX Wash Buffer and blocked with EnVision FLEX Peroxidase Block for 5 min. Primary antibody (Ki67 (Cat. No. M7240, RRID:AB_2142367, dilution 1:170, Agilent, Santa Clara, CA) or cleaved caspase-3 (Cat. No. 9664, RRID:AB_2070042, dilution 1:600, Cell Signaling Technology, Danvers, MA)) incubation was performed at an optimized concentration and duration, followed by extensive washing. Sections were then incubated with FLEX/HRP-labeled polymer for 30 min, washed, and developed using DAB+ Substrate-Chromogen for 5 min. Counterstaining was performed with Biocare CAT Hematoxylin for 5 s, followed by bluing for 30 s. Finally, sections were washed, dehydrated, cleared, and cover-slipped for microscopic analysis. Images were acquired using Zeiss Axio Vert.A1 inverted microscope under 10x magnification.

### Statistical Analysis

For analysis of the means of three or more groups, analysis of variance (ANOVA) tests were performed. In the event that ANOVA justified post-hoc comparisons between group means, the comparisons were conducted using Tukey’s multiple-comparisons test. A two-sided unpaired Student’s *t*-test was used for comparisons between the means of two groups. *P* <0.05 was considered to be statistically significant.

### Data Availability

All the data related to this manuscript are available in main figures, supplemental information, or upon reasonable request from the corresponding author.

## RESULTS

### A combination of DT2216 and everolimus potentiates anti-tumor activity of MRTX1133 *in vitro* by apoptosis induction

We performed long-term colony formation assays to test the combinations of DT2216/everolimus with MRTX1133 in AsPC1, HPAC, SW1990 and PANC1 cells. All three compounds as single agent had significant, but partial inhibition of colony growth. MRTX1133 alone reduced only 40-60% cell colonies compared to DMSO-control. The addition of DT2216 or everolimus to MRTX1133 significantly reduced colony formation compared to MRTX1133 alone. More importantly, the triple combination of MRTX1133+DT2216+everolimus was significantly potent than any of the two-drug combinations and leads to near complete inhibition of colony growth in AsPC1, HPAC, and SW1990 cell lines (**Fig. 1a-c; e-g**). Though, the triple combination had a less pronounced effect in PANC1 cells compared to the other three cell lines as about 25% colonies were able to survive compared to control, but the effect was significant compared to single agents or two-drug combinations (**Fig. 1d & h**).

**Figure 1.**
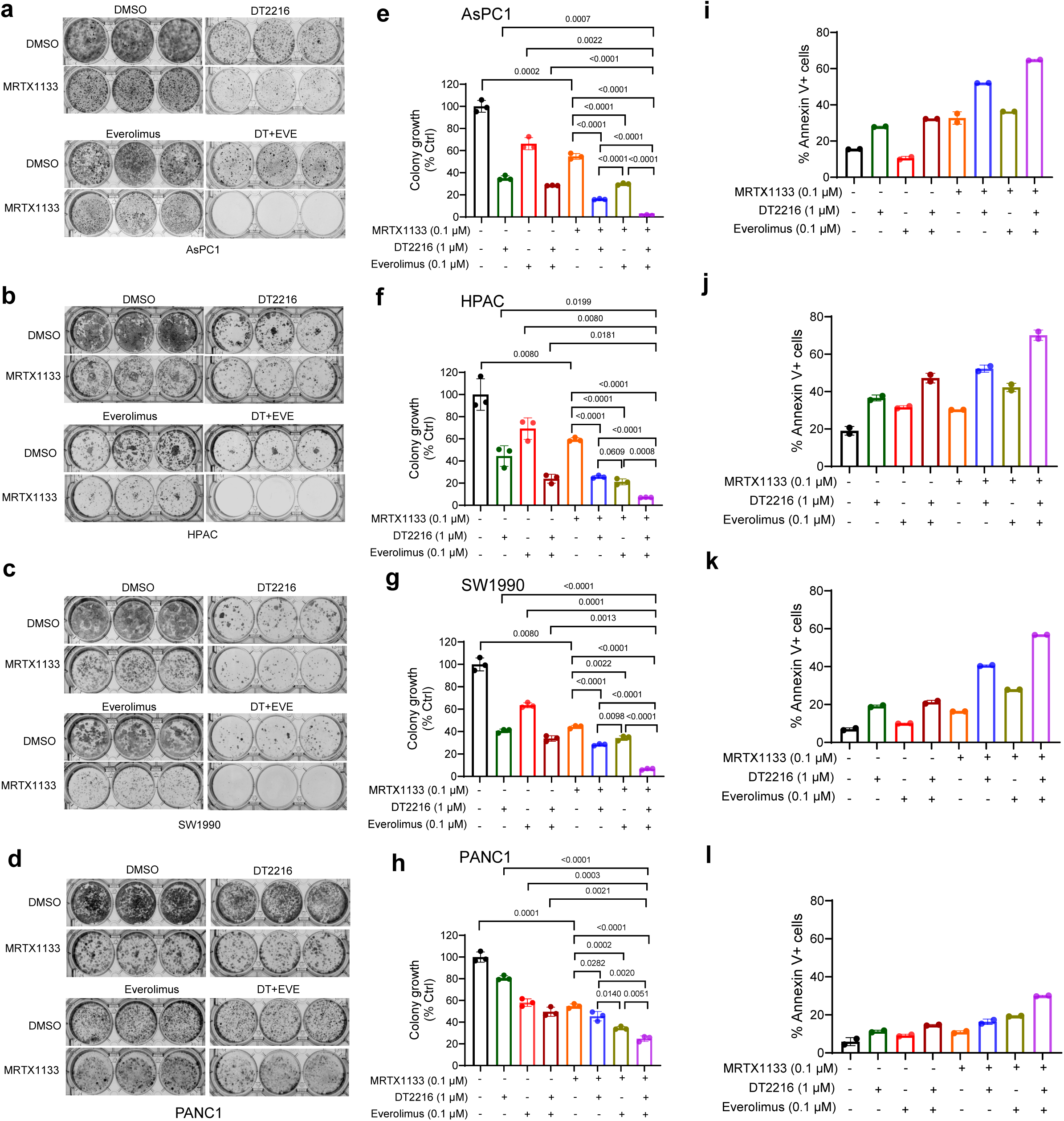
A combination of DT2216 and everolimus enhances anti-clonogenic effect of MRTX1133 through promoting apoptosis. a-d. Colony images of AsPC1 (a), HPAC (b), SW1990 (c) and PANC1 cells (d) after they were treated with MRTX1133, DT2216, everolimus and their combinations as indicated for 10-14 days followed by crystal violet staining. **e-h.** Colorimetric measurement of colony growth in AsPC1 (e), HPAC (f), SW1990 (g) and PANC1 cells (h). Data are presented as mean ± SD (n =3 cell culture replicates). Statistical significance was determined by two-sided Student’s t-test, and *p* values are mentioned in the graphs. **i-l.** Percentage annexin V^+^ apoptotic population in AsPC1 (i), HPAC (j), SW1990 (k) and PANC1 cells (l) after they were treated with MRTX1133, DT2216, everolimus and their combinations at indicated concentrations of for 48 h. Data are presented as mean ± SD (n =2 independent experiments).

Next, we performed annexin V/PI staining followed by flow cytometric analysis for quantitative measurement of apoptosis. The results indicate modest apoptosis induction by MRTX1133 alone, which was enhanced when it was combined with DT2216 or everolimus, and the highest apoptosis induction was obtained with the triple combination (**Fig. 1 i-l**). The results from apoptosis analysis were also correlated with those obtained with colony formation assays. In addition, we measured cellular apoptosis by immunoblotting analysis of cleaved caspase-3 (CC3) and cleaved PARP. MRTX1133 alone induced minimal apoptosis as indicated by barely detectable CC3 and cleaved PARP levels. However, the addition of DT2216 leads to substantial increase of both CC3 and cleaved PARP, which was further increased after the triple combination treatment (**Suppl. Fig. 1a-c**). Overall, these results confirm that MRTX1133 leads to enhanced tumor cell killing by apoptosis when combined with DT2216 and everolimus.

### MRTX1133 increases pro-apoptotic BH3-only protein BIM and decreases NOXA in G12D-mutated PDAC cells

In our previous study, we found that G12Ci sotorasib increases BIM expression in G12C-mutated tumor cell lines.^18^ Therefore, we wondered whether MRTX1133 can also induce BIM expression in G12D-mutated PDAC cells. Consistent with our hypothesis, we found that MRTX1133 can substantially increase BIM expression in multiple G12D-mutated PDAC cell lines including both p53-mutated (AsPC1, HPAC, and PANC1) and p53 wild-type (SW1990) (**Fig. 2a-h**). However, the extent of BIM upregulation after MRTX1133 treatment was cell line-dependent with highest upregulation in SW1990 cells and least upregulation in PANC1 cells. In addition, the expression of NOXA – a pro-apoptotic protein that endogenously binds to and inhibits anti-apoptotic MCL-1 – was substantially reduced after MRTX1133 treatment in all these cell lines, with most potent reduction in PANC1 cells (**Fig. 2a-h**). However, MRTX1133 treatment exerted no observable effects on expressions of BCL-xL and MCL-1 (**Suppl. Fig. 2a-d**). Interestingly, SW1990 cells also showed the highest KRAS inhibition, as indicated by decreased phosphorylated p-ERK1/2, and p-Akt compared to other cell lines. The least BIM upregulation in PANC1 cells is correlated with its lesser sensitivity to KRAS inhibition.^12, 30, 31^ In addition, MRTX1133 treatment leads to compensatory increase in p-AKT reflecting an activation of the PI3K/AKT survival pathway, which can provide an alternative route for cell survival and proliferation, potentially reducing sensitivity of PANC1 cells to MRTX1133. Additionally, minimal inhibition of p-S6 suggests incomplete suppression of mTOR signaling, further contributing to reduced MRTX1133 in PANC1 cells. In HPAC cells, p-AKT and p-S6 inhibitions were observed at relatively higher doses of MRTX1133. We confirmed that MRTX1133-mediated BIM upregulation and NOXA downregulation was specific to G12D-mutated PDAC cells, as MRTX1133 exerted no considerable effect on their expressions in KRAS wild-type BxPC3 cells (**Suppl. Fig. 3a & b**). Also, MRTX1133 treatment exerted no observable effects on expressions of BCL-xL and MCL-1 in BxPC3 cells (**Suppl. Fig. 3c**).

**Figure 2.**
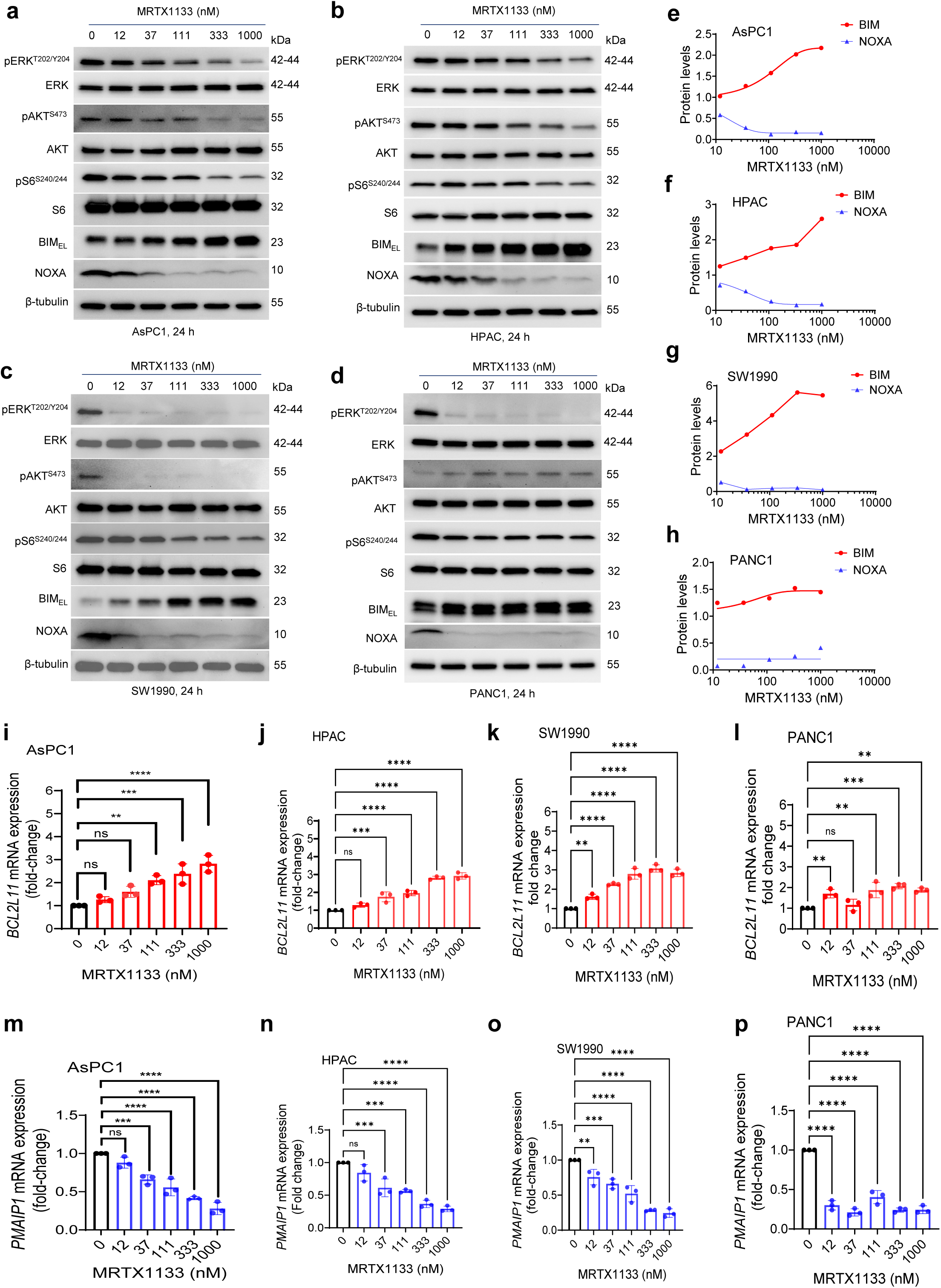
MRTX1133 increases BIM and suppresses NOXA expression in G12D-mutated PDAC cells. a-d. Immunoblot analyses of phosphorylated- and total- ERK, AKT and S6, BIM extra-long isoform (BIM_EL_) and NOXA in AsPC1 (a), HPAC (b), SW1990 (c), and PANC1 cells (d) after they were treated with indicated concentrations of MRTX1133 for 24 h. The β-tubulin was used as an equal loading control. **e-h.** Densitometric analysis of BIM_EL_ and NOXA immunoblots in AsPC1 (e), HPAC (f), SW1990 (g), and PANC1 cells (h). Immunoblots detect three isoforms of BIM i.e., short isoform (BIM_S_), long isoform (BIM_L_) and extra-long isoform (BIM_EL_). Among them, BIM_EL_ is the major isoform and is shown here. **i-l.** mRNA expression levels of BIM-coding gene *BCL2L11* in AsPC1 (i), HPAC (j), SW1990 (k), and PANC1 cells (l) after 24 h treatment with MRTX1133. **m-p**. mRNA expression levels of NOXA-coding gene *PMAIP1* in AsPC1 (m), HPAC (n), SW1990 (o), and PANC1 cells (p) after 24 h treatment with MRTX1133. Data in **i-p** are presented as mean ± SD (n= 3 biological replicates). \*\**p* <0.01, \*\*\**p* <0.001, \*\*\*\**p* <0.0001, ^ns^ *p* >0.05 i.e., not significant compared to untreated (control) cells as determined by one-way ANOVA and Dunnett’s multiple comparisons test. Since ERK1 and ERK2 have very close molecular weights (∼44 and ∼42 kDa), the two bands appeared as a single merged band in panels a-d (and subsequent figures) due to limited resolution in gradient gels that we have used.

Next, we evaluated whether MRTX1133 can modulate mRNA levels of genes encoding BIM and NOXA in these PDAC cell lines. We found a dose-dependent and significant increase in mRNA levels of *BCL2L11* (BIM-coding gene) and reduction in *PMAIP1* (NOXA-coding gene), which were correlated with the protein levels after MRTX1133 treatment in different cell lines (**Fig. 2i-p**). For example, the upregulation of *BCL2L11* was least potent and downregulation of *PMAIP1* was most potent in PANC1 cells (**Fig. 2l & p**).

### MRTX1133-mediated increased BIM is sequestered by BCL-xL and is released by DT2216, while reduced NOXA is partially compensated by everolimus to inhibit MCL-1

In order to elucidate the mechanisms of synergistic anti-tumor effect of MRTX1133 when combined with DT2216 and everolimus, we first determined the effects of the DT2216/everolimus combination with MRTX1133 on KRAS signaling, and expressions of BCL-xL, BIM and NOXA. The triple combination leads to enhanced inhibition of p-ERK and p-S6 coupled with BCL-xL degradation, BIM upregulation and restoration of NOXA (**Fig. 3a**).

**Figure 3.**
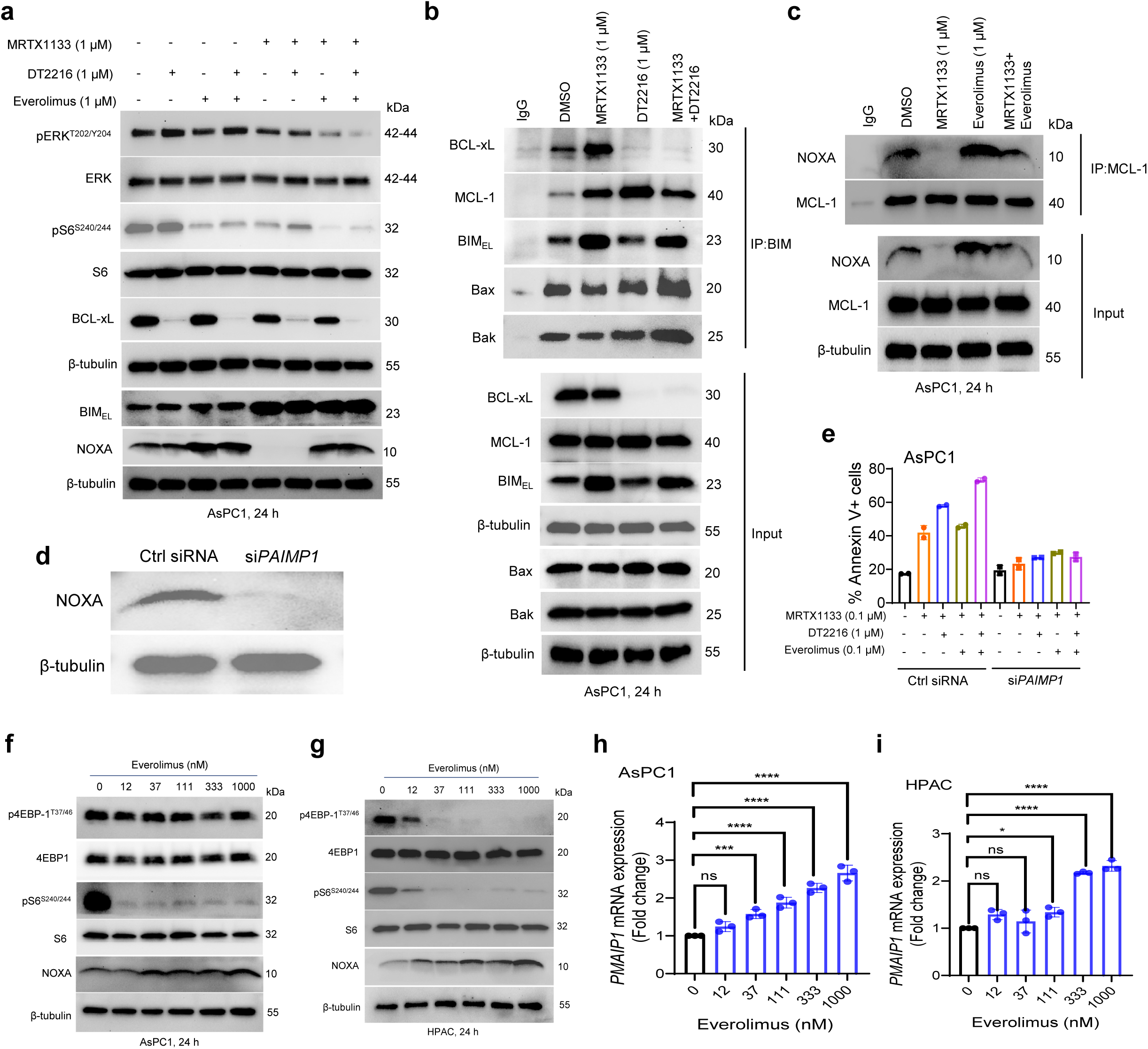
Everolimus-mediated mTOR inhibition increases NOXA and DT2216-mediated BCL-xL degradation frees BIM. **a**. Immunoblot analyses of phosphorylated- and total- ERK and S6, BCL-xL, BIM and NOXA in AsPC1 cells after they were treated with MRTX1133, DT2216, everolimus and their combinations as indicated for 24 h. **b**. Immunoprecipitation analysis of BIM in AsPC1 cells after they were treated with DMSO, MRTX1133, DT2216 or MRTX1133+DT2216 for 24 h, and the immunoprecipitated as well as input samples were subjected to immunoblot analysis of BCL-X_L_, MCL-1, and BIM. **c**. Immunoprecipitation analysis of MCL-1 in AsPC1 cells after they were treated with DMSO, MRTX1133, everolimus or MRTX1133+everolimus for 24 h, and the immunoprecipitated as well as input samples were subjected to immunoblot analysis of NOXA and MCL-1. **d, e.** siRNA-mediated *PMAIP1* (gene encoding NOXA) knockdown in AsPC1 cells as confirmed by immunoblotting (d), and percentage annexin V^+^ apoptotic population in Control (Ctrl) siRNA and *PAIMP1* siRNA exposed AsPC1 cells after they were treated with MRTX1133 and its combinations with DT2216 and/or everolimus at indicated concentrations for 48 h (e). Data in **e** are presented as mean ± SD (n = 2 independent experiments). **f, g.** Immunoblot analyses of phosphorylated- and total- 4EBP1 and S6, and NOXA in AsPC1 (f), and HPAC cells (g) after they were treated with indicated concentrations of everolimus for 24 h. **h, i**. mRNA expression levels of NOXA-coding gene *PMAIP1* in AsPC1 (h) and HPAC cells (i) after 24 h treatment with MRTX1133. Data are presented as mean ± SD (n= 3 biological replicates). \**p* <0.05, \*\*\**p* <0.001, \*\*\*\**p* <0.0001, ^ns^ *p* >0.05 i.e., not significant compared to untreated (control) cells as determined by one-way ANOVA and Dunnett’s multiple comparisons test. β-tubulin was used as an equal loading control in all immunoblots.

In our previous publication, we have reported that G12Ci-induced BIM is sequestered by BCL-xL, and therefore BIM is not free to induce apoptosis in G12C-mutated tumor cells.^18^ Here, we wondered whether MRTX1133-induced BIM is also sequestered by BCL-xL, and whether DT2216-mediated BCL-xL degradation can release BIM to induce apoptosis. Our immunoprecipitation (IP) analysis showed that BCL-xL binds to MRTX1133-induced BIM in AsPC1 cells. Further treatment with DT2216 caused BCL-xL degradation and released BIM. BIM has propensity to also bind with MCL-1; therefore, we immunoblotted IP:BIM samples with an MCL-1 antibody. It turned out that MCL-1 can bind to BIM, although with lesser intensity than BCL-xL. Furthermore, MCL-1 can bind to a portion of MRTX1133-induced BIM, and upon DT2216 treatment, slightly more BIM is available to bind MCL-1 because of unavailability of BCL-xL. However, with MRTX1133/DT2216 combination treatment, BIM binding to MCL-1 was slightly reduced compared to DT2216 alone, suggesting that an extra amount of free BIM is available to activate BAX and BAK and ultimately apoptosis. To further investigate this, we performed immunoblotting analysis of BAX and BAK on IP:BIM samples. These results show that after combination treatment with MRTX1133+DT2216, increased BIM binds to BAX and BAK (**Fig. 3b**). Thereby, BIM facilitates BAX/BAK activation to induce mitochondrial apoptosis. In another experiment, we immunoprecipitated MCL-1 and immunoblotted these samples with a NOXA antibody in order to determine whether increased NOXA by everolimus can promote MCL-1/NOXA complex formation. We found that MRTX1133 treatment leads to diminished NOXA/MCL-1 complex formation, while co-treatment with everolimus leads to partial restoration of NOXA/MCL-1 complex formation (**Fig. 3c**). The binding of NOXA to MCL-1 leads to MCL-1 inhibition, thus enabling apoptosis induction.^32, 33^ We extended this mechanistic validation to HPAC (relatively sensitive to MRTX1133) and PANC1 (relatively resistant to MRTX1133) cells. In HPAC cells, the results were similar to those observed in AsPC1 (**Suppl. Fig. 4a–c**). In PANC1 cells, on the other hand, the triple combination leads to inhibition of pERK, but stabilization of NOXA was not obvious. Further, there was only minimal binding of BIM to BCL-xL or MCL-1 in MRTX1133-treated PANC1 cells. Similarly, there was almost negligible binding between NOXA and MCL-1 (**Supplementary Fig. 4d-f**).

To further confirm the functional role of NOXA in mediating apoptosis, we performed siRNA-mediated knockdown of *PMAIP1* (the gene encoding NOXA) in AsPC1 cells and assessed apoptosis using Annexin V/PI staining and flow cytometry following treatment with MRTX1133 alone and in combination with everolimus ± DT2216, compared to control siRNA-exposed cells. *PMAIP1* knockdown remarkably diminished apoptosis induction by MRTX1133/everolimus ± DT2216 combinations (**Fig. 3d & e**). These findings provide direct evidence that NOXA induction by everolimus is required for robust apoptosis in combination therapy.

Furthermore, we confirmed that everolimus leads to a dose-dependent upregulation of NOXA at both protein and mRNA levels in G12D-mutated AsPC1 and HPAC cell lines (**Fig. 3f-i**). However, unlike our and others published reports that mTOR inhibition causes MCL-1 suppression in different tumor cells,^26, 34, 35^ everolimus did not alter the expression of MCL-1 in these G12D-mutated PDAC cell lines (**Suppl. Fig. 4g-i**). These results suggest that a combination of DT2216 and everolimus can potentially enhance the anti-tumor activity of MRTX1133 through BIM release and NOXA-mediated MCL-1 inhibition, respectively, and potentially in part, through enhanced inhibition of ERK and mTOR.

We finally evaluated the effect of direct targeting MCL-1 using the selective inhibitor S63845 in KRAS G12D-mutated AsPC1 cells, both as a single agent and in combination with MRTX1133 and/or DT2216. These results indicate that the combination of S63845 with MRTX1133 significantly increased cell loss compared to MRTX1133 alone, and the triple combination of MRTX1133, S63845, DT2216 was significantly more potent than single agents and two drug combinations (**Supplementary Fig. 5a-c**). However, S63845 in combination with MRTX1133 and/or DT2216 exerts slightly less significant antitumor effects as seen with MRTX1133/everolimus or MRTX1133/DT22216/everolimus combinations in G12D-mutated AsPC1 tumor cells (**Fig. 1a & e**).

### The DT2216/everolimus combination enhances the efficacy of MRTX1133 *in vivo*

We evaluated the antitumor efficacy of the combination of MRTX1133 with DT2216/everolimus in vivo using AsPC1 xenograft model. As anticipated, DT2216 alone had no significant effect on tumor growth. MRTX1133 and everolimus alone leads to minimal tumor growth inhibition, but the combination of DT2216 or everolimus with MRTX1133 leads to significant tumor growth inhibition compared to individual agents. Importantly, the triple combination leads to the greatest tumor inhibition, which was significant compared to all other groups. More importantly, the triple combination caused tumor regressions in five out of seven mice in the group (**Fig. 4a-b**). Neither the two-drug combinations nor the triple combination leads to any changes in mouse body weights (**Fig. 4c**). In addition, the triple combination did not show increased toxicity to mice based on blood cell counts (platelets, WBCs and RBCs) (**Suppl. Fig. 6a-c**), and liver/kidney function as measured by ALT/AST activities (**Suppl. Fig. 6d & e**). Of note, DT2216 alone caused some platelet reduction, but the levels remained above 0.1 million per µL of blood, which is considered clinically safe.^18, 24, 26, 36^ Moreover, the addition of everolimus and/or MRTX1133 did not cause further reduction in platelet counts. These results suggest that the triple combination of MRTX1133 with DT2216 and everolimus is more efficacious compared to the two-drug combinations.

**Figure 4.**
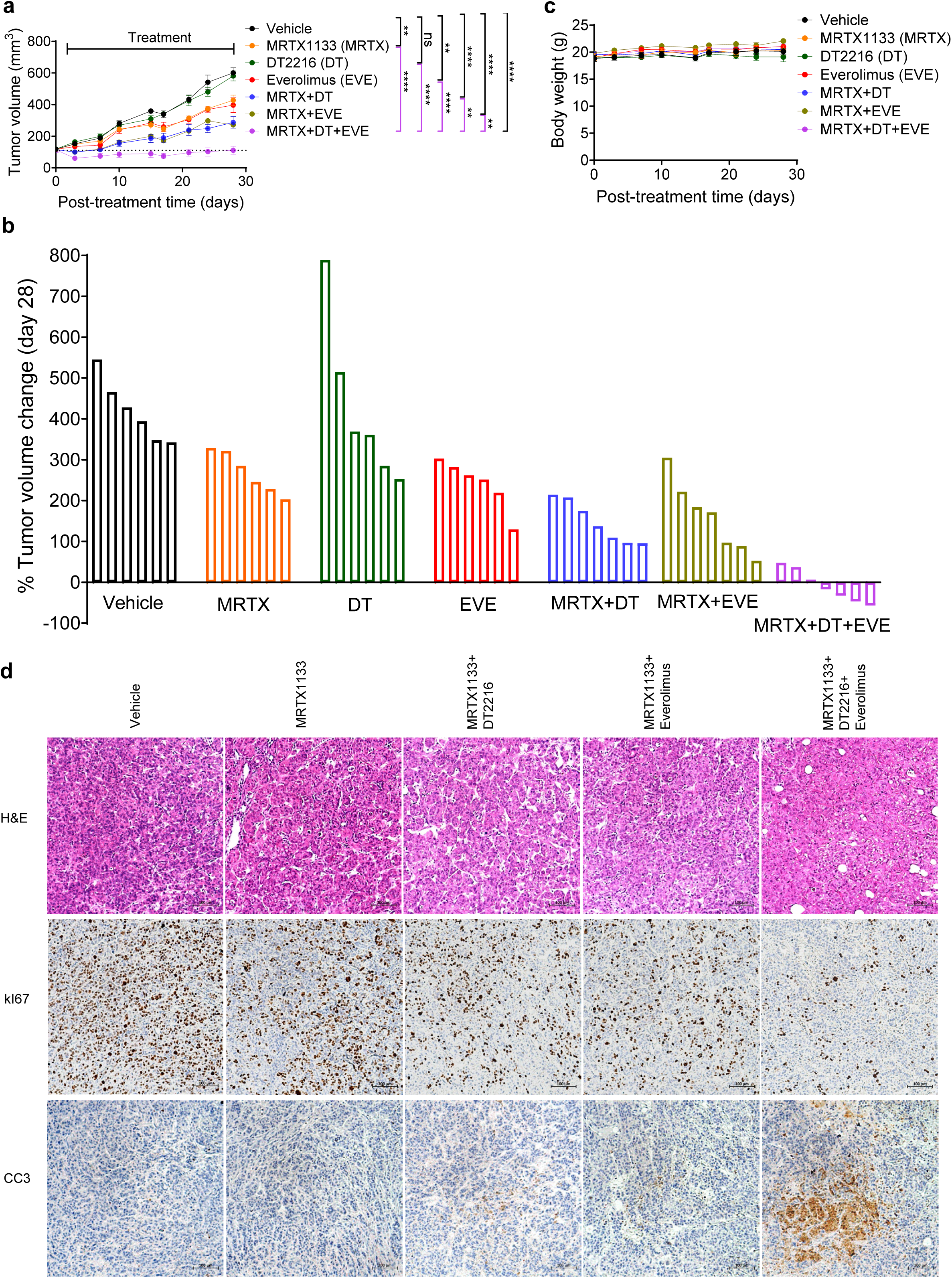
The DT2216/everolimus combination enhances the efficacy of MRTX1133 in AsPC1 xenograft model. **a**. Tumor volume changes in AsPC1 xenografts after they were treated with vehicle, MRTX1133 (MRTX, 3 mg/kg, b.i.d, 5 days ON/2 days OFF, i.p.), DT2216 (DT, 15 mg/kg, 2x/week, i.p.), everolimus (EVE, 2.5 mg/kg, 2x/week, p.o.) or their combinations as indicated. Data are presented as mean ± SEM (n = 6-7 mice per group). Statistical significance was determined by two-sided Student’s *t*-test at the last tumor measurement i.e., day 28. ** *p* <0.01, **** *p* <0.0001, ^ns^ *p* >0.05 i.e., not significant. **b**. Percent changes in tumor volumes of individual mice at the end of treatment as compared to their baseline tumor volumes. **c**. Mouse body weight changes after treatment as in **a**. **d,** H&E, Ki67 and CC3 staining in tumor samples from PD study.

We next measured PD markers of MRTX1133, DT2216 and everolimus after short-term treatment of AsPC1 xenografts in a separate experiment. DT2216 leads to a substantial reduction in BCL-xL levels which sustained in two-drug and triple combination groups. MRTX1133 at the used dosage of 3 mg/kg bid leads to a substantial reduction of p-ERK, but not of p-S6 and p-AKT. While S6 phosphorylation was effectively reduced after everolimus treatment and sustained in combination treated tumors. We did not see any further inhibition of p-ERK or p-S6 over that of MRTX1133 or everolimus, respectively, in the combination groups (**Suppl. Fig. 7a-b**). This suggests the triple combination exerts enhanced antitumor efficacy by simultaneously targeting KRAS, BCL-xL and mTOR. H&E staining of tumor tissues showed that the vehicle group exhibited large, densely packed tumor cells with areas of necrosis. Treatment with MRTX1133 alone moderately reduced tumor size and cellularity, but tumor cells remained substantial. The combination of MRTX1133 with DT2216 or everolimus showed further tumor shrinkage and increased apoptosis. The triple combination of MRTX1133 with DT2216 and everolimus was the most effective, resulting in the smallest tumors with minimal tumor cellularity and increased necrosis, indicating an enhanced antitumor effect (**Fig. 4d**). IHC analysis of Ki-67 expression demonstrated high levels of staining in the vehicle group. Treatment with MRTX1133 reduced Ki67^+^ cells moderately, which were further reduced by its combination with DT2216 or everolimus. Importantly, the triple combination leads to the best effect in reducing Ki67^+^ cells. On the other hand, an opposite trend was observed with CC3 staining, with highest staining in the triple combination group (**Fig. 4d**). These results suggest that the triple combination leads to enhanced antitumor activity by simultaneous inhibition of cell proliferation and induction of apoptosis.

### The DT2216/everolimus combination can overcome acquired MRTX1133 resistance both in vitro *and* in vivo

Acquired resistance is a key factor contributing to the limited clinical efficacy of KRAS inhibitors. We generated MRTX1133 acquired-resistant AsPC1 cells (designated AsPC1-MR) by exposing them to progressively increasing concentrations of MRTX1133. The AsPC1-MR cells showed a substantial increase in resistance to MRTX1133 compared to parental AsPC1 cells (**Fig. 5a**). Thereafter, we performed immunoblotting analysis on AsPC1-MR cells and their parental AsPC1 counterparts to assess the expression of selected BCL-2 family proteins. While no noticeable changes were observed in BCL-xL and BCL-2 levels, a substantial increase in BIM, along with a decrease in NOXA, was detected in AsPC1-MR cells compared to parental cells. In addition, AsPC1-MR cells also showed a marked increase in MCL-1 levels (**Fig. 5b**). Next, we investigated whether the combination of DT2216 and everolimus could sensitize AsPC1-MR cells to MRTX1133 treatment. While DT2216 or everolimus alone moderately sensitized AsPC1-MR cells to MRTX1133, the DT2216+everolimus combination markedly enhanced their sensitivity in both short-term cell viability and long-term colony formation assays (**Fig. 5c-h**). Of note, the acquired resistance to MRTX1133 was associated with increased sensitivity to everolimus, suggesting that cellular dependence on mTOR pathway increases upon development of resistance to MRTX1133, consistent with a previous report.^12^ (**Fig. 5c-e**). This suggests that mTOR plays a more critical role in the acquired resistance of AsPC1-MR cells to MRTX1133, whereas BCL-xL has predominant role in intrinsic resistance. Indeed, AsPC1-MR cells exhibited a marked increase in mTOR activation, as evidenced by elevated p-S6 levels. Additionally, increased phosphorylation of ERK and AKT was also observed in AsPC1-MR cells (**Fig. 5i**). Furthermore, we explored the mechanism underlying the sensitizing effect of the DT2216+everolimus combination and found that the combination simultaneously inactivates p-ERK and p-S6, degrades BCL-xL, and restores NOXA expression in AsPC1-MR cells (**Fig. 5j**). Interestingly, everolimus was found to slightly decrease BIM expression in AsPC1-MR cells; however, the remaining BIM levels appeared sufficient to induce cell killing.

**Figure 5.**
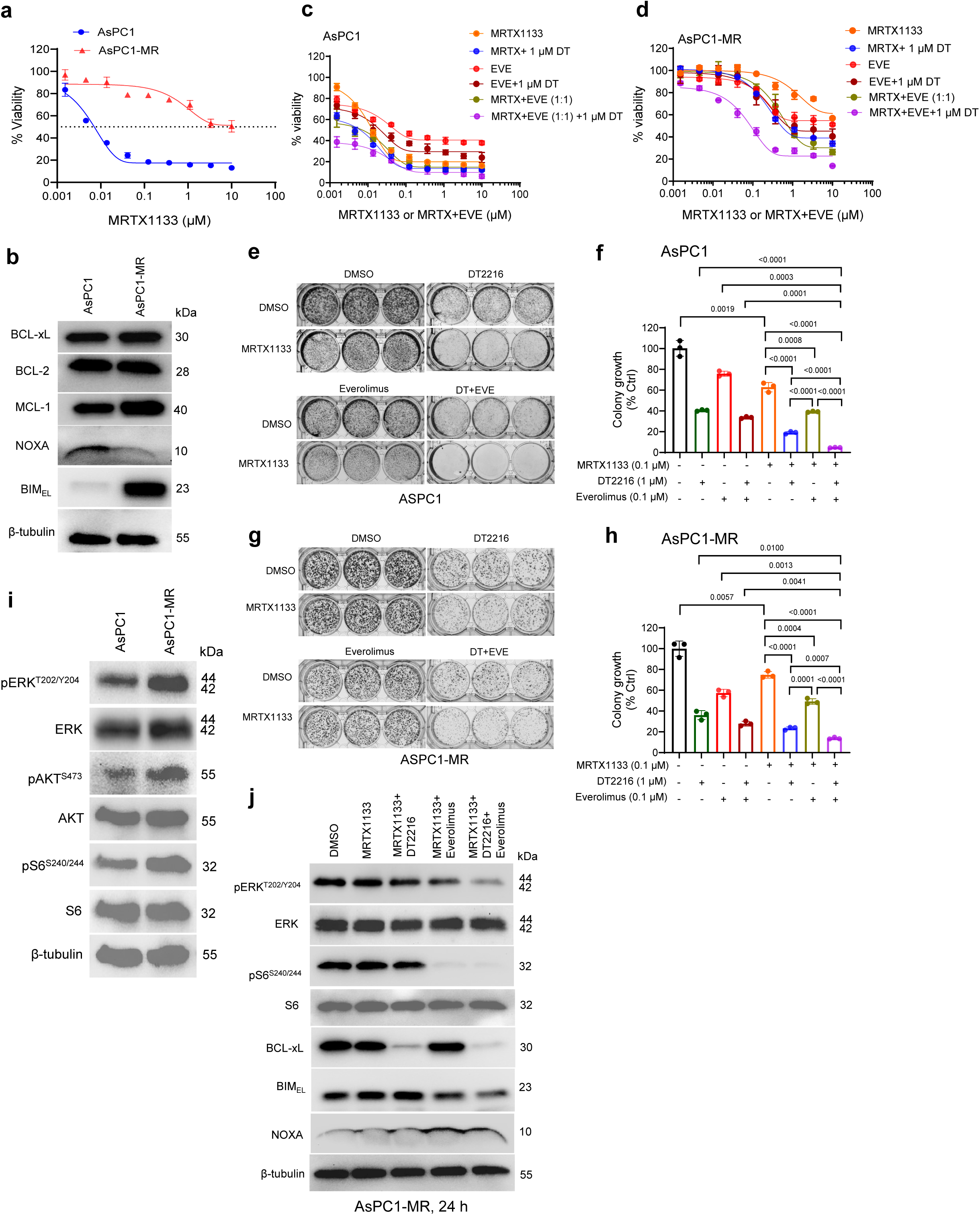
The DT2216/everolimus combination sensitizes resistant cells to MRTX1133 treatment *in vitro*. **a.** Viability of MRTX1133-resistant AsPC1 (AsPC1-MR) cells compared to parental AsPC1 cells after they were treated with increasing concentrations of MRTX1133 for 72 h. **b**. Immunoblot analyses of BCL-xL, BCL-2, MCL-1, NOXA and BIM in AsPC1 and AsPC1-MR cells. **c, d**. Viability of parental AsPC1 (c) and AsPC1-MR cells (d) after they were treated with increasing concentrations of MRTX1133, everolimus (EVE) alone and/or with 1 µM of DT2216 (DT) and/or equimolar (1:1) ratio of everolimus for 72 h. Data are presented as ± SD (n = 3 replicate cell cultures). First points in MRTX + 1 µM DT2216, EVE + 1 µM DT2216 and MRTX + EVE (1:1) + 1 µM DT2216 represent 1 µM DT2216 alone. **e, g**. Colony images of parental AsPC1 (e) and AsPC1-MR cells (g) after they were treated with MRTX1133, DT2216 or EVE alone or with dual or triple combinations as indicated for 14 days followed by crystal violet staining. **f, h**. Colorimetric measurement of colony growth in parental AsPC1 (f) and AsPC1-MR cells (h). Data in **a**, **c, d, f** and **h** are presented as mean ± SD (n = 3 cell culture replicates). Statistical significance for data in panels **f** and **h** was determined by two-sided Student’s t-test, and *p* values are mentioned in the graphs. **i**. Immunoblot analyses of phosphorylated- and total- ERK, AKT and S6 in AsPC1 and AsPC1-MR cells. **j**. Immunoblot analyses of phosphorylated- and total- ERK, S6, and AKT, BCL-xL, BIM and NOXA in AsPC1-MR cells after they were treated with MRTX1133 alone or in combination with DT2216 and/or everolimus as indicated. β-tubulin was used as an equal loading control in all immunoblots.

We next generated an AsPC1-CDX-MR model by treating AsPC1 tumor-bearing mice (average tumor size 161 mm^3^) with MRTX1133 at 10 mg/kg twice a day (bid). Initially, mice responded to treatment with continued tumor regressions and average tumor reached 66 mm^3^ after 17 days of treatment, thereafter tumors started regrowing and grown to 146 mm^3^ after 28 days. At this point, tumors were considered resistant for our study. Thereafter, mice were randomized into four groups and treated with MRTX1133 alone, MRTX1133 + DT2216, MRTX1133 + everolimus, or MRTX1133 + DT2216 + everolimus (**Fig. 6a**). As expected, MRTX1133-treated mice exhibited continuous tumor growth. MRTX1133 + DT2216-treated mice exhibited tumor regression for one week before tumors began to regrow. MRTX1133 + everolimus-treated mice showed tumor inhibition for approximately two weeks, after which tumors started growing again. MRTX1133 + DT2216 + everolimus-treated mice demonstrated sustained tumor regression for up to four weeks before tumors began to regrow, albeit more slowly than in the other treatment groups. Notably, one mouse in the MRTX1133 + DT2216 + everolimus group had a significantly larger tumor than the other mice; however, this tumor showed substantial regression following the triple combination treatment (**Fig. 6b**). None of the treatment groups showed changes in body weight during or at the end of the treatment (**Fig. 6c**). The tumor growth inhibition in these mice was associated with BCL-xL degradation by DT2216, reduction of pERK by MRTX1133+everolimus, and reduction of pS6 by everolimus in resistant tumors. Moreover, the triple combination led to enhanced inhibition of p-ERK and p-S6 (**Suppl. Fig. 8a & b**).

**Figure 6.**
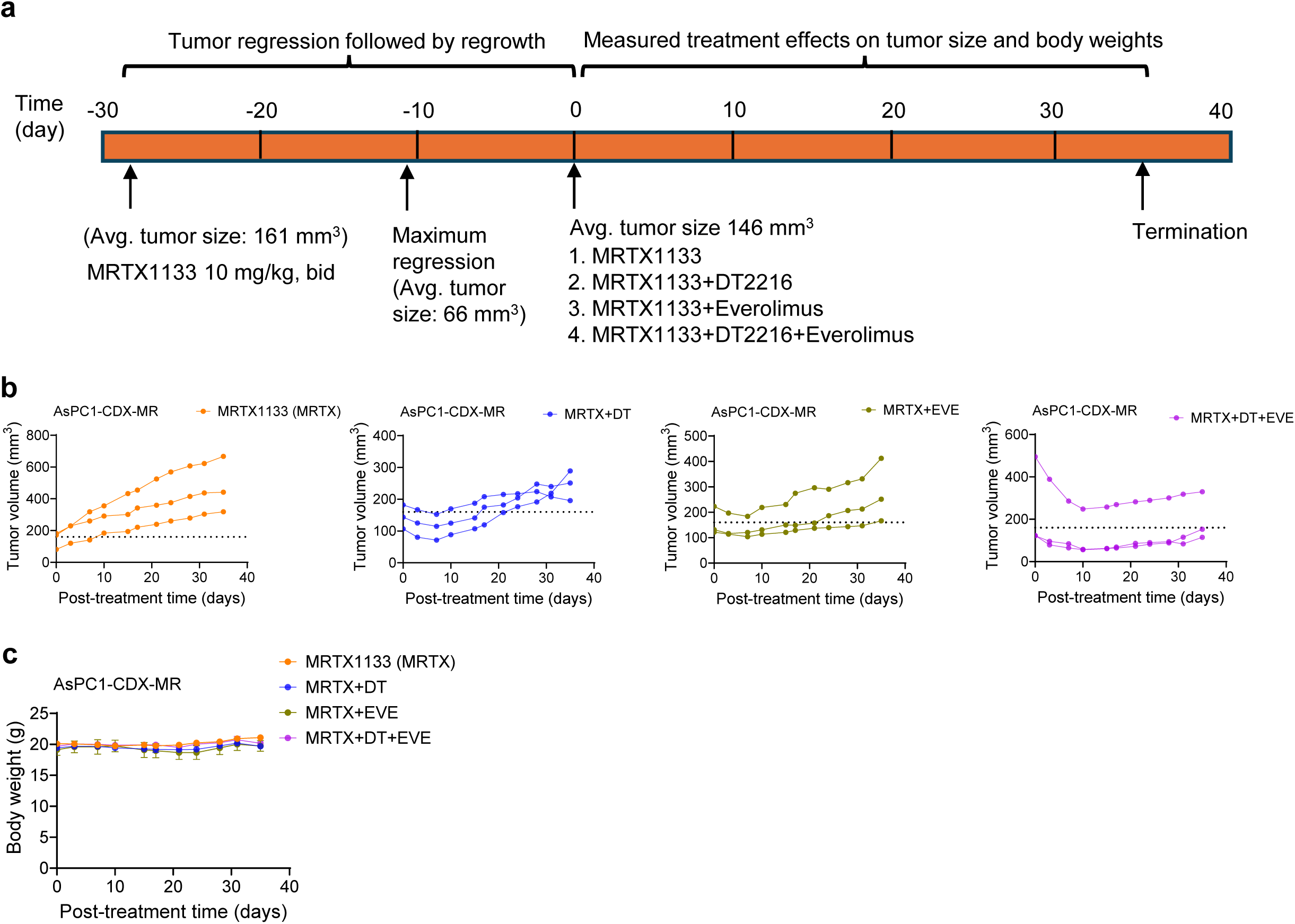
The DT2216/everolimus combination sensitizes resistant AsPC1 xenografts to MRTX1133 treatment. **a.** Schematic of AsPC1-CDX-MR study design. **b**. Tumor volume changes in individual AsPC1-CDX-MR mice after they were treated with MRTX1133 (MRTX, 10 mg/kg, b.i.d, i.p.) alone or its combination with DT2216 (DT, 15 mg/kg, 2x/week, i.p.) and/or everolimus (EVE, 2.5 mg/kg, 2x/week, p.o.). **c**. Mouse body weight changes during the course of treatment as in **b.**

## DISCUSSION

Based on our previous study and those of others, G12Ci’s have been shown to induce minimal apoptosis, due to a dysregulated expression of BCL-2 family proteins, contributing to both intrinsic and acquired resistance.^18, 19^ In the current study, we primarily focus on intrinsic resistance mechanisms to G12Di MRTX1133 within G12D-mutant PDAC tumors. Specifically, resistance to MRTX1133 appears to be driven by impaired apoptosis (e.g., due to overexpression of anti-apoptotic proteins such as BCL-xL and MCL-1) rather than alternate pathway activation alone as reported previously.^12–14, 37^ We observed that MRTX1133 can only partially inhibit the colony forming capabilities of multiple G12D-mutated tumor cells with mild to modest apoptosis induction. The triple combination of DT2216 and everolimus with MRTX1133 was found to dramatically eliminate colony growth and was significantly more potent than the two-drug combinations of DT2216/MRTX1133 or everolimus/MRTX1133. Although the triple combination was significantly more potent than the MRTX1133 or two-drug combinations in PANC1 cells, it could not completely eliminate clonogenic growth, suggesting PANC1 cells are outlier among G12D-mutated tumor cells. Of note, the two-drug combinations of DT2216/MRTX1133 and everolimus/MRTX1133 showed heterogeneous effects in different tumor cells. DT2216/MRTX1133 combination was more potent in AsPC1 and SW1990 cells, whereas everolimus/MRTX1133 was more potent in PANC1 cells, and they were similar in HPAC cells. Though MRTX1133 alone could only induce mild to modest apoptosis, its combination with DT2216 or DT2216/everolimus robustly induced apoptosis. These results confirm ours and others previous findings that KRAS inhibitors generally act as cytostatic agents with only minimal apoptosis induction as a single agent and therefore may require combinations with apoptosis inducers for optimum therapeutic efficacy.^18, 19^ This is because the impaired expression of anti-apoptotic proteins, particularly BCL-xL and MCL-1, in PDAC cells limit the ability of MRTX1133 monotherapy to induce cell death, even when KRAS signaling is effectively suppressed.^18, 28, 38–40^

Furthermore, we demonstrated that G12Di MRTX1133 leads to an abolishment of KRAS signaling predominantly through inhibition of the ERK/MAPK pathway specifically in G12D-mutated PDAC cells. Notably, the PI3K/Akt/mTOR pathway was differentially inhibited by MRTX1133 in different G12D-mutated PDAC cell lines, and was not significantly inhibited in PANC1 cells, which have previously been reported to relatively resistant to KRAS signaling inhibitors.^12, 30, 31^ Since KRAS regulates both ERK/MAPK and PI3K/Akt pathways, both pERK and pAkt are regarded as biomarkers of KRAS inhibition.^4, 6^ However, our data indicate that MRTX1133-responsive cells (AsPC1, HPAC and SW1990) exhibit more pronounced decreases in pAkt compared to resistant cells (PANC1), which maintain PI3K/Akt signaling despite KRAS inhibition. Therefore, our data suggest that pAkt inhibition could be better predicter of response to MRTX1133. Importantly, we observed a significant upregulation of BIM and downregulation of NOXA following MRTX1133 treatment specifically in the G12D-mutated PDAC cell lines, whereas expressions of BCL-xL and MCL-1 remained unaffected.

Apoptosis is regulated by the balance between pro-apoptotic proteins (such as BIM and NOXA) and anti-apoptotic proteins (such as BCL-xL and MCL-1).^41, 42^ By modulating the expression of BIM and NOXA, MRTX1133 can induce a dependence on BCL-xL and MCL-1 in G12D-mutated PDAC cells. BCL-xL and MCL-1 are overexpressed and play a critical role in promoting survival in many solid tumors, including PDAC.^38–40^ The increased levels of BIM can be sequestered by high amounts of BCL-xL, preventing BIM from freely activating BAK/BAX, an essential step for intrinsic or mitochondrial-mediated apoptosis induction.^38, 41, 42^ Additionally, the reduced expression of NOXA may unleash MCL-1, which further binds to BIM, contributing to a survival advantage for tumor cells.^41^ Since NOXA is a pro-apoptotic BH3-only protein that endogenously binds to and inhibits MCL-1,^43–45^ the reduction of NOXA by MRTX1133 can potentially contribute to apoptosis resistance through an increased activity of MCL-1. Therefore, strategies to restore NOXA expression may have potential to enhance the efficacy of MRTX1133 by promoting apoptosis. Based on this, we hypothesize that BCL-xL degradation in combination with inhibiting MCL-1 via NOXA restoration could be a potential mechanism of enhanced efficacy of MRTX1133. BCL-xL degradation can be achieved with the PROTAC DT2216 without affecting platelets, unlike conventional BCL-xL inhibitors.^24^ However, available MCL-1 inhibitors lack tumor selectivity and can induce cardiotoxicity and hepatotoxicity, particularly when combined with BCL-xL inhibitors or degraders.^46, 47^ Therefore, MCL-1 inhibition via NOXA could be a clinically relevant strategy. The mTOR inhibitors have been shown to suppress MCL-1 expression in various tumor cells, as demonstrated by us and others.^26, 34, 35^ To our surprise, the mTOR inhibitor everolimus could not reduce MCL-1 levels in G12D-mutated PDAC cells, but it notably upregulated NOXA expression. This could be partly because mTOR inhibition is known to trigger a variety of cellular stress responses, including inhibition of cap-dependent translation and induction of the integrated stress response (ISR).^48, 49^ It has been reported that under certain stress conditions, NOXA can be upregulated via ATF4, a key transcription factor activated during ISR. Therefore, mTOR inhibition may increase NOXA expression through activation of the ATF4 pathway, leading to transcriptional upregulation of *PMAIP1*, the gene encoding NOXA.^50, 51^ Further, Immunoprecipitation analysis revealed that the increased NOXA following everolimus treatment binds to MCL-1, potentially inhibiting its function. However, everolimus can enhance apoptosis not only through partial restoration of NOXA but also through inhibition of mTOR signaling, because feedback activation of mTOR signaling is known to contribute to resistance to KRAS inhibitors.^13, 52^ In addition, we confirmed the requirement for NOXA in the induction of apoptosis by MRTX1133/everolimus ± DT2216 combinations by genetic knockdown of *PMAIP1* (the gene encoding NOXA) in AsPC1 cells. Additionally, DT2216 induced BCL-xL degradation, thereby increasing free BIM levels to activate BAK/BAX.

*In vivo*, we found that MRTX1133 alone or its combination with DT2216 or everolimus are able to slow down tumor growth. However, the triple combination not only completely inhibited tumor growth, but also induced tumor regressions that sustained through the end of treatment in a majority of the tumor bearing mice. Moreover, we did not observe any significant changes in mouse body weights, or increased toxicity to blood cells or impaired liver function suggesting the triple combination is tolerable in mice. Notably, DT2216 as single agent had no significant antitumor effect despite BCL-xL degradation and inhibiting colony growth in vitro. The lack of single-agent efficacy of DT2216 in vivo is likely due to compensatory survival signals present in the tumor microenvironment, and/or a shift in survival reliance on other anti-apoptotic proteins like MCL-1 upon BCL-xL depletion. Therefore, BCL-xL loss alone is insufficient to exert an anti-tumor effect unless accompanied by pro-apoptotic signals, such as BIM upregulation, which is achieved with MRTX1133 co-treatment. Through PD analysis, we found that MRTX1133, DT2216 and everolimus leads to inhibition/degradation of their respective targets, suggesting on-target antitumor activity. In addition, the triple combination leads to more pronounced inhibition of Ki67 and induction of cleaved caspase-3 in tumors after short-term treatment. Therefore, tumor regressive activity of the triple combination was likely attributed to simultaneous inhibition of tumor cell proliferation and enhanced apoptosis. However, in tumors, we did not see enhanced inhibition of p-ERK or p-S6 unlike in vitro results, which might be because these mice were only treated for five days.

We next established MRTX1133 acquired resistant AsPC1 cells (AsPC1-MR) and xenograft model. In AsPC1-MR cells, we found a decreased expression of NOXA, and increased BIM compared to parental AsPC1 cells. In addition, these cells showed increased activation of ERK, AKT and mTOR. The triple combination of MRTX1133, DT2216 and everolimus significantly inhibited colony growth compared to MRTX1133 alone or two-drug combinations in AsPC1-MR cells. Interestingly, everolimus was slightly more effective in AsPC1-MR cells as compared to parental AsPC1 cells. This might have resulted from increased dependence of AsPC1-MR cells on mTOR pathway, as indicated by higher mTOR activity, compared to parental AsPC1 cells consistent with a published report.^12^ We also found an increased inhibition of ERK after triple combination treatment in AsPC1-MR cells, suggesting that the increased ERK activity was suppressed. In AsPC1 xenograft model that became resistant after continuous treatment with MRTX1133, the triple combination leads to sustained tumor regressions for prolonged duration than two-drug combinations. However, we hypothesize that MRTX1133-resistant cells harboring secondary KRAS mutations—particularly those that directly affect MRTX1133 binding—are unlikely to respond to the combination of everolimus and DT2216, as MRTX1133 would no longer be effective as a backbone agent. Our combination approach is intended to enhance the apoptotic response in MRTX1133-sensitive or partially resistant cells, but it is unlikely to overcome resistance driven by secondary KRAS mutations. However, this combination strategy may delay or reduce the emergence of acquired resistance by inducing robust apoptosis and preventing survival of residual tumor cells.

In addition to apoptotic dysregulation, EMT has been reported as a key driver of resistance to KRASi in PDAC, particularly in PANC-1 cells, which exhibit a mesenchymal phenotype associated with reduced drug sensitivity. EMT-mediated plasticity enables tumor cells to evade KRAS-targeted therapy by activating alternative survival pathways and promoting invasive characteristics.^12–14^ While our study did not directly assess EMT, the limited response observed in PANC-1 cells may partly reflect EMT-related resistance, underscoring the importance of future investigations into strategies that combine KRASi with agents targeting EMT or its downstream signaling.

For our studies, we used 3 mg/kg twice daily and 10 mg/kg twice daily dosing of MRTX1133 for AsPC1 xenograft and acquired resistant AsPC1 xenograft, respectively, which are significantly lower than maximum tolerated dose (MTD) of 30 mg/kg twice daily. For the resistance model (Fig. 6), we increased the dose of MRTX1133 to 10 mg/kg twice daily dosing based on results at 3 mg/kg twice daily dosing (Fig. 4) that did not induce tumor regression as a single agent and tumors continued to grow from the outset. Since our goal in the resistance model was to first achieve tumor regression followed by regrowth, we selected the higher dose. Moreover, we also used a lower dose of everolimus (2.5 mg/kg) at lower frequency (twice a week). These combinations were well tolerated in mice without any significant toxicities, as measured by body weights, blood cell counts and AST/ALT enzymatic activities for liver/kidney function. Since the higher doses of MRTX1133 (upto 30 mg/kg, i.e., MTD) have been used in different combination studies,^4, 53, 54^ future studies will evaluate if the increasing dosing of MRTX1133 and/or everolimus can lead to complete responses in patient-derived xenograft models of G12D-mutated PDAC without increasing normal toxicities.

### LIMITATIONS OF THE STUDY

The siRNA-mediated genetic knockdown of NOXA gene suggests NOXA’s involvement in apoptosis induction, however the lack of studies involving complete knockout of NOXA gene by CRISPR-Cas9 system and in vivo validation is lacking that would have further strengthened these findings. Our current in vivo studies are limited to a relatively sensitive AsPC1 xenograft model, therefore further validation in a relatively resistance cell line, such as PANC1, would be necessary to establish whether the combination therapy can sensitize these intrinsically resistant tumors. Since desmoplastic stroma may be an important resistance mechanism to MRTX1133 in PDAC patients, our study has limitation of not using immunocompetent mouse model (such as orthotopic syngeneic models and genetically engineered mouse models (GEMMs)) to more closely recapitulate human disease. In addition, for our acquired resistance model, we used only three mice per group with the primary goal of assessing resistance and sensitization to MRTX1133. Considering a small cohort of mice used, these studies are preliminary and require further validation with larger cohorts and alternative models (e.g., transplantation of resistant tumors in new mice) to more rigorously investigate the combination effect on MRTX1133 acquired resistance. Furthermore, these studies are limited by a lack of IHC staining of apoptosis markers, such as TUNEL or cleaved caspase-3, in resistant tumors.

## CONCLUSION

Our study provides strong evidence that the combination of DT2216 and everolimus enhances the efficacy of MRTX1133 by potentiating apoptosis induction. Using both *in vitro* and *in vivo* models, including those with acquired resistance, we demonstrated that this combination significantly enhances therapeutic efficacy compared to MRTX1133 alone through increased apoptotic cell death and reduced tumor growth. These findings highlight the potential of this combination therapy in overcoming resistance mechanisms in G12D-mutated PDAC. After further evaluation in PDX models and spontaneous tumor models (such as KPC model) of G12D-mutated PDAC, this approach could pave the way for future clinical exploration of this strategy to improve G12Di efficacy in patients.

## ACKNOWLEDGEMENTS

This study was supported in parts by the US National Institutes of Health (NIH) grant (R01 CA242003) to D. Zhou, G. Zheng & J. Trevino; William and Ella Owens Medical Foundation Grant to D. Zhou & S. Khan; and Mike-Hogg Fund to S. Khan. Funding from the American Cancer Society (ACS) Pilot Grant and the Mays Cancer Center (MCC) support grant to S. Khan is gratefully acknowledged. We are thankful to the Biospecimen and Translational Genomics Core at the UT Health San Antonio for their help with H&E and IHC analyses.

## Notes

### Competing Interest Statement

P. Zhang , G. Zheng, D. Zhou and S. Khan are inventors of patent applications for the use of BCL-xL PROTACs (including DT2216) as antitumor agents. G. Zheng and D. Zhou are co-founders of and have equity in Dialectic Therapeutics, which develops BCL-xL PROTACs to treat cancer.

