## Supplementary Figures S1-S7; Supplementary Table S1 for "The combination of BCL-xL PROTAC and mTOR inhibitor sensitizes pancreatic ductal adenocarcinoma to KRAS^G12D^ inhibitor treatment by enhancing apoptosis induction"

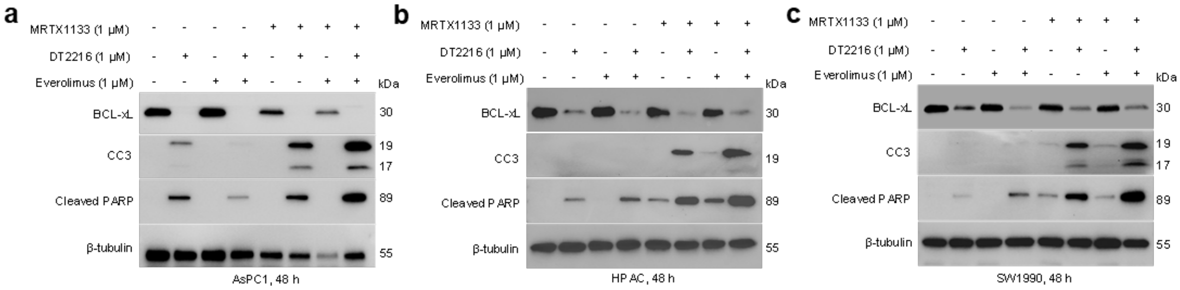


**Suppl. Fig. 1**. **a-c.** Immunoblot analyses of BCL-xL, cleaved caspase-3 (CC3) and cleaved PARP in AsPC1 (a), HPAC (b) and SW1990 cells (c) after they were treated with MRTX1133, DT2216, everolimus and their combinations as indicated for 48 h. β-tubulin was used as an equal loading control in all immunoblots.

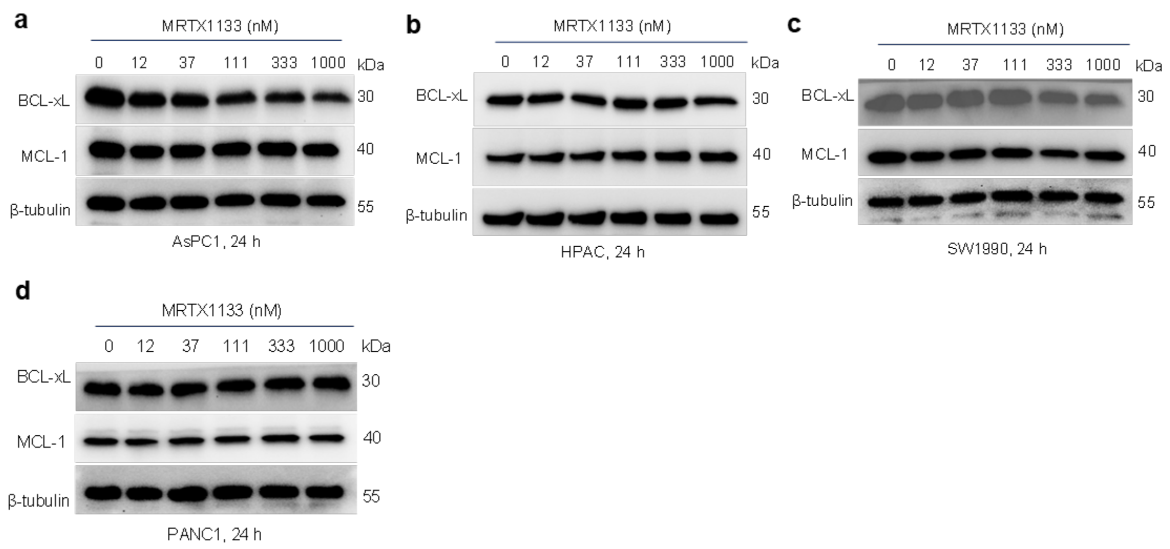


**Suppl. Fig. 2**. **a-d**. Immunoblot analyses of BCL-xL and MCL-1 in AsPC1 (a), HPAC (b), SW1990 (c), and PANC1 (d) cells after they were treated with indicated concentrations of MRTX1133 for 24 h. The β-tubulin was used as an equal loading control.

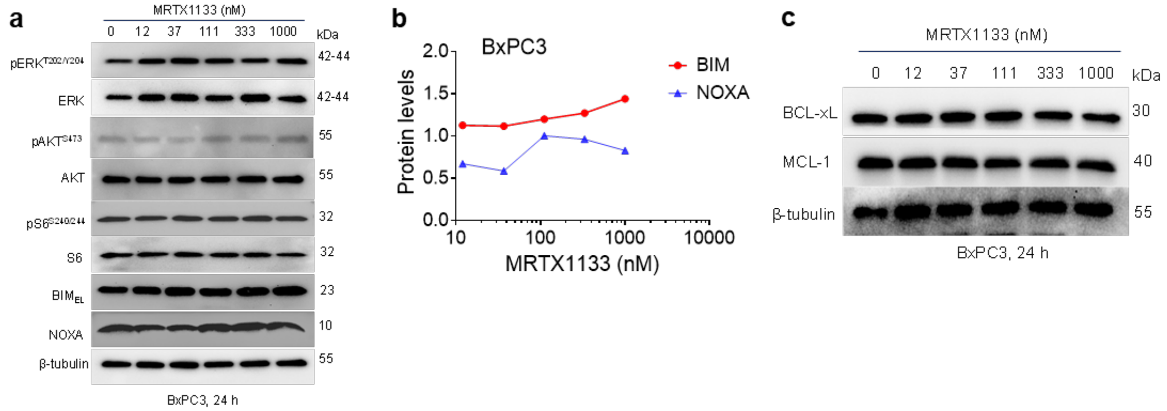


**Suppl. Fig. 3**. **a.** Immunoblot analyses of phosphorylated- and total- ERK, AKT and S6, BIM extra-long isoform ($BIM_{EL}$) and NOXA in AsPC1 cells after they were treated with indicated concentrations of MRTX1133 for 24 h. **b.** Densitometric analysis of $BIM_{EL}$ and NOXA immunoblots in AsPC1 cells. **c**. Immunoblot analyses of BCL-xL and MCL-1 in BxPC3 cells. The β-tubulin was used as an equal loading control.

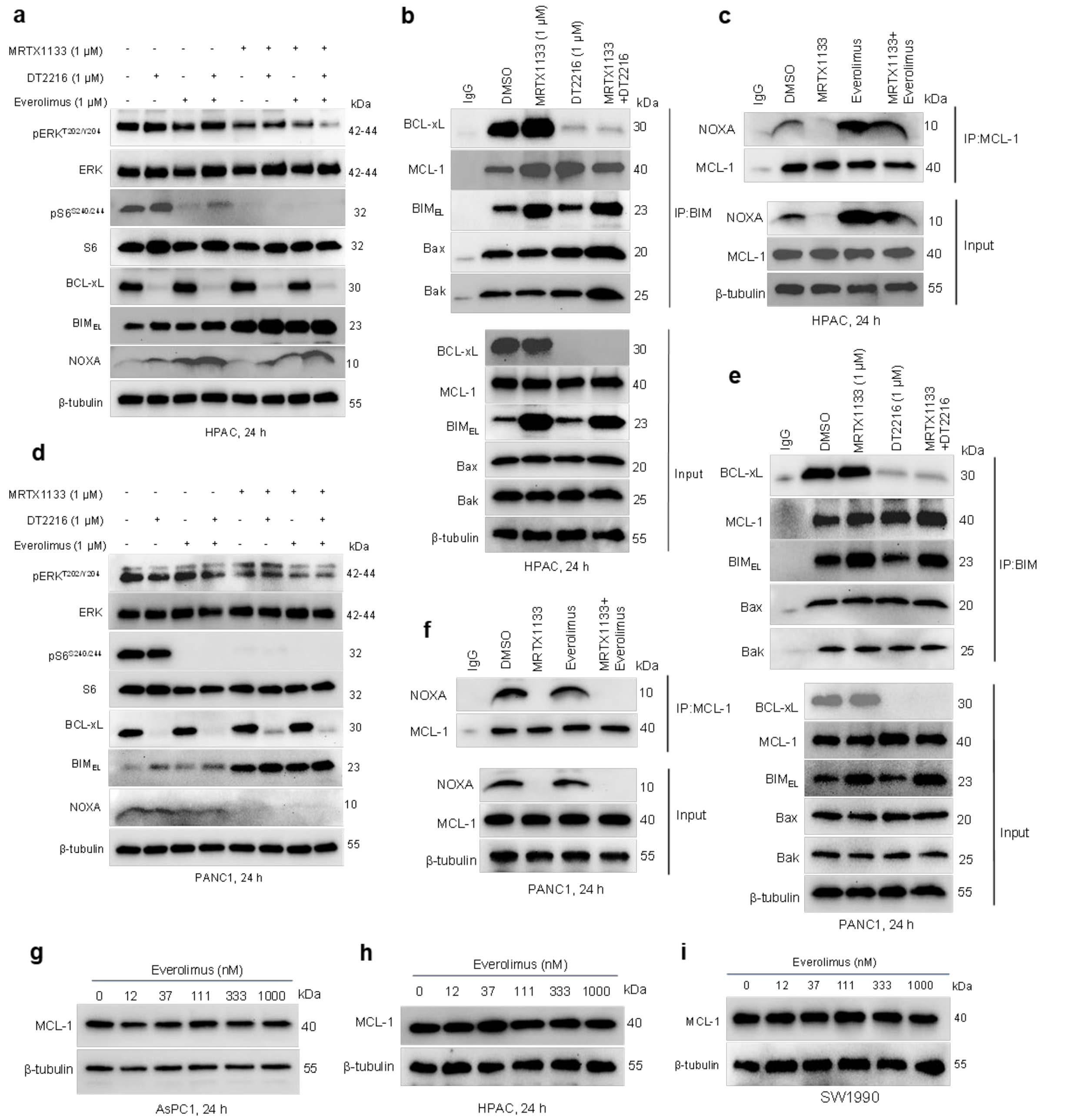


**Supplementary Figure S4. a-c. a, d**. Immunoblot analyses of phosphorylated- and total-ERK and S6, BCL-xL, BIM and NOXA in HPAC (a) and PANC1 (d) cells after they were treated with MRTX1133, DT2216, everolimus and their combinations as indicated for 24 h. **b, e**. Immunoprecipitation analysis of BIM in HPAC (b) and PANC1 (e) cells after they were treated with DMSO, MRTX1133, DT2216 or MRTX1133+DT2216 for 24 h, and the immunoprecipitated as well as input samples were subjected to immunoblot analysis of BCL-$X_L$, MCL-1, and BIM. **c, f**. Immunoprecipitation analysis of MCL-1 in HPAC (c) and PANC1 (f) cells after they were treated with DMSO, MRTX1133, everolimus or MRTX1133+everolimus for 24 h, and the immunoprecipitated as well as input samples were subjected to immunoblot analysis of NOXA and MCL-1. **g-i.** Immunoblot analyses of MCL-1 in AsPC1 (g), HPAC (h) and SW1990 cells (i) after they were treated with indicated concentrations of everolimus for 24 h. The β-tubulin was used as an equal loading control.

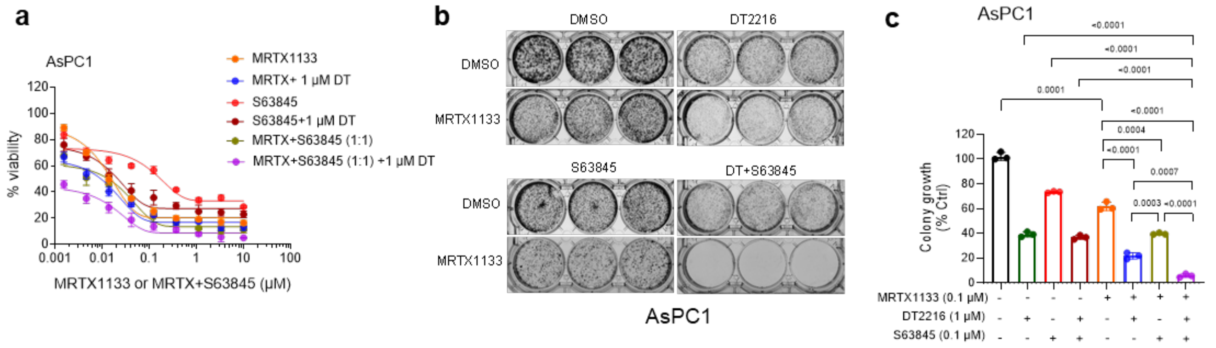


**Suppl. Fig. 5. a.** Viability of parental AsPC1 cells after they were treated with increasing concentrations of MRTX1133, S63845 alone and/or with 1 µM of DT2216 and/or equimolar (1:1) ratio of S63845 for 72 h. Data are presented as ± SD (n= 3 replicate cell cultures). First points in MRTX + 1 µM DT2216, S63845 + 1 µM DT2216 and MRTX + S63845 (1:1) + 1 µM DT2216 represent 1 µM DT2216 alone. **b.** Colony images of AsPC1 cells after they were treated with MRTX1133, DT2216, S63845 and their combinations as indicated for 14 days followed by crystal violet staining. **c.** Colorimetric measurement of colony growth in AsPC1 cells. Data are presented as mean ± SD (n =3 cell culture replicates). Statistical significance was determined by two-sided Student's t-test, and *p* values are mentioned in the graphs.

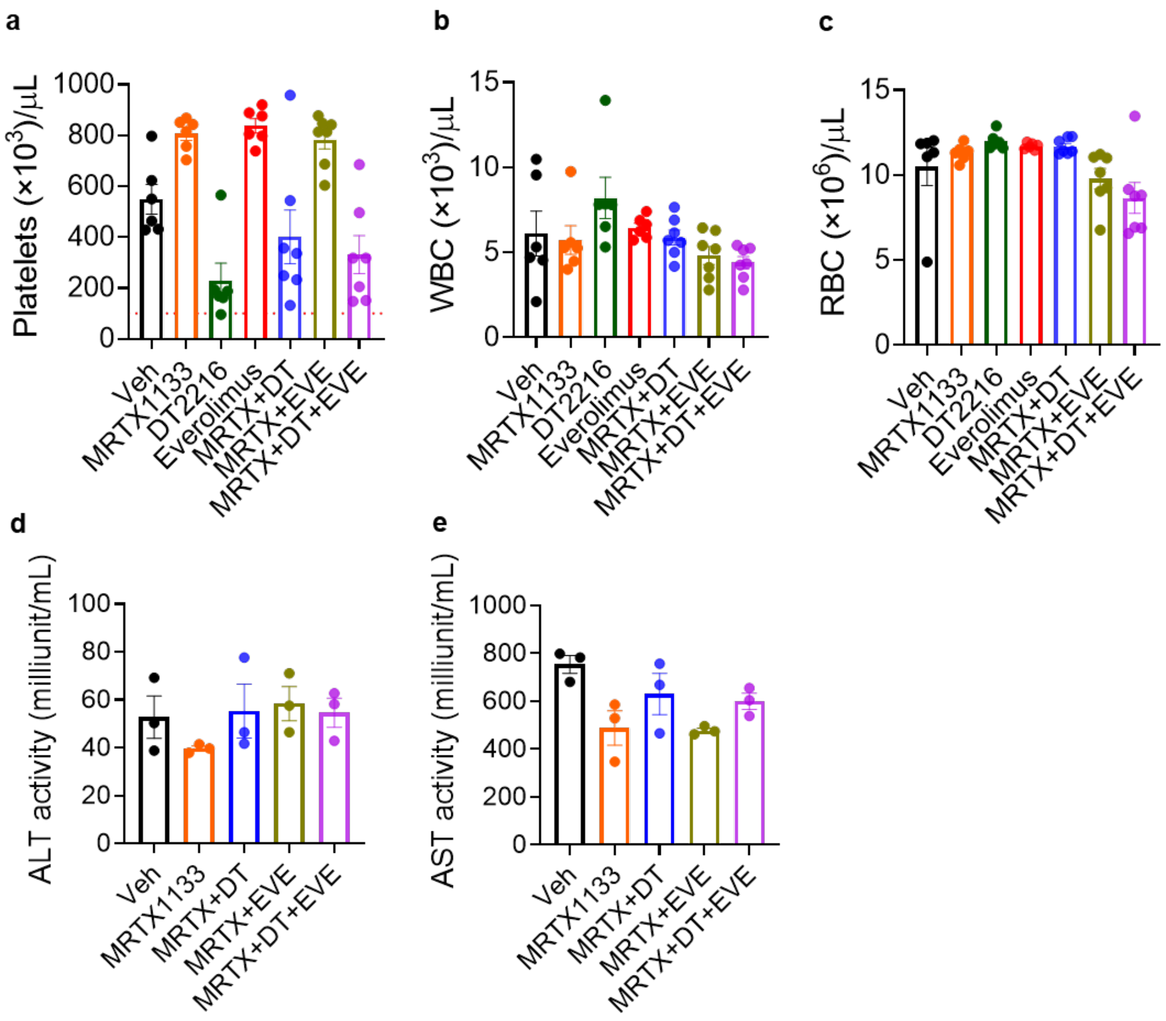


**Suppl. Fig. 6. a-c.** Enumeration of platelets (a), WBCs (b), and RBCs (c) 24 h after the final dose of DT2216. Data are presented as mean ± SEM (n = 6-7 mice per group). **d, e**. ALT (d) and AST (e) activities in mouse serum as measured 24 h after final dose of DT2216. Data are presented as mean ± SEM (n = 3 mice per group).

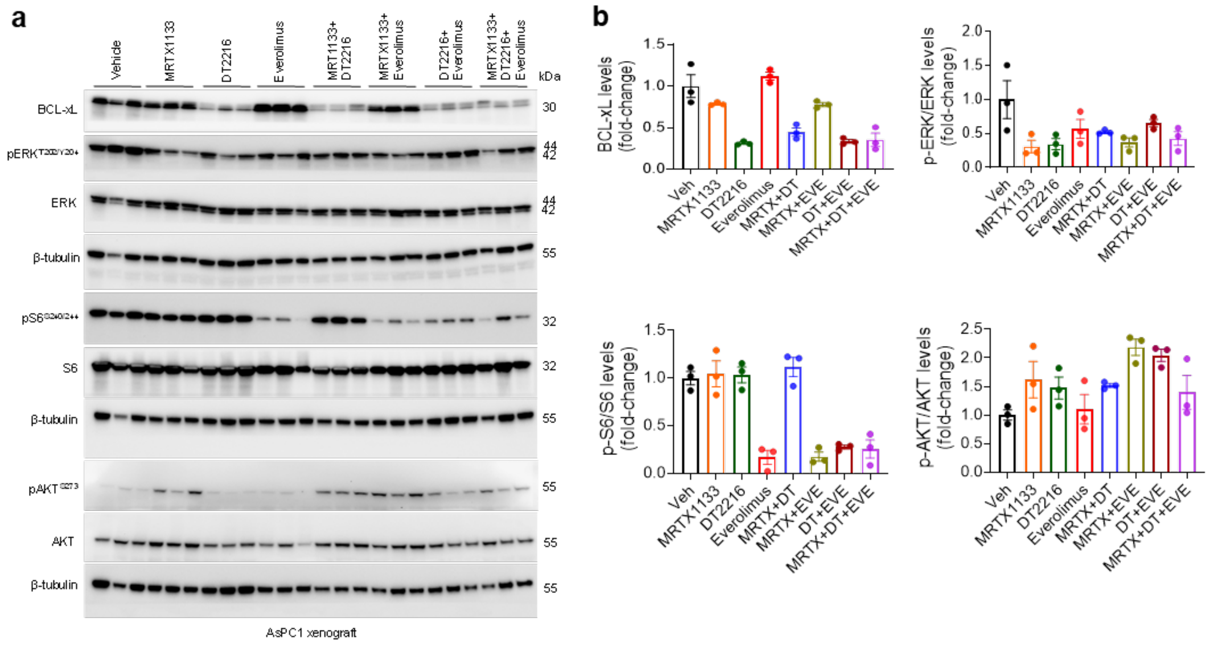

**Suppl. Fig. 7. The MRTX1133, DT2216 and everolimus demonstrate their respective target inhibition in AsPC1 xenografts. a.** Immunoblot analyses of BCL-xL, phosphorylated- and total- ERK, S6, and AKT in AsPC1 xenograft mice after they were treated with vehicle, MRTX1133 (MRTX, 3 mg/kg, b.i.d, Day 1-5, i.p.), DT2216 (DT, 15 mg/kg, Day 1 and Day 5, i.p.), everolimus (EVE, 2.5 mg/kg, Day 1 and Day 5, p.o.) or their combinations as indicated and tumors were harvested 24 h after 2$^{nd}$ dose of DT2216 i.e., Day 6. The β-tubulin was used as an equal loading control. **b.** Densitometric analysis of immunoblots as in **a**. Data are presented as mean ± SEM (n = 3 mice per group).

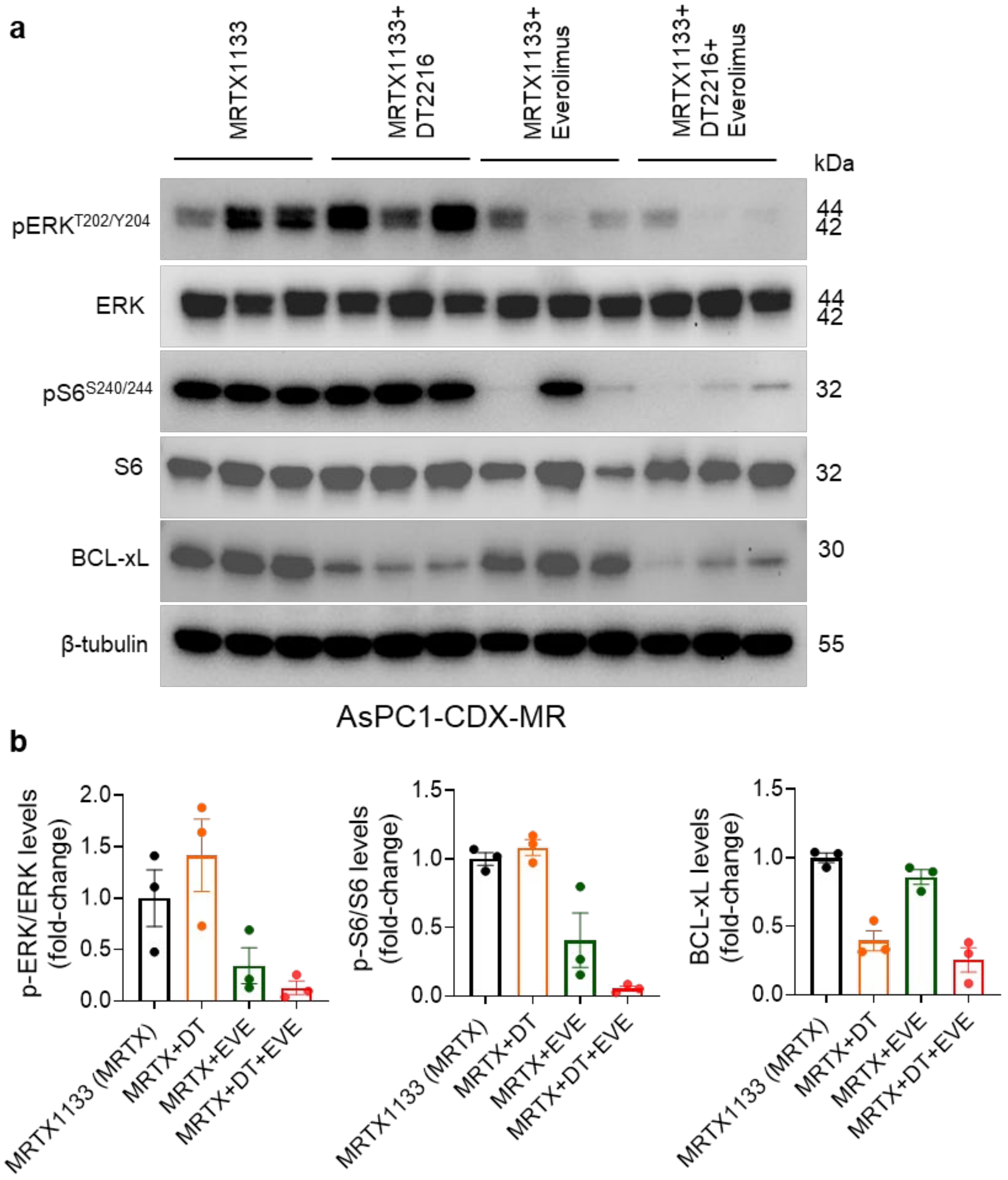


**Suppl. Fig. 8. a.** Immunoblot analyses of phosphorylated- and total- ERK and S6, BCL-xL, BIM and NOXA in AsPC1-CDX-MR tumors after indicated treatments. **b**. Densitometric analysis of immunoblots as in **a**. Data are presented as mean ± SEM (n = 3 mice per group).

**Supplementary Table S1.** Antibodies used in immunoblotting.

| Antibody | Clone | Antibody | Catalog | RRID | Concentration |
|---|---|---|---|---|---|
| p-ERK T202/Y204 | _ | Rabbit IgG | 9101 | AB_331646 | 1:1000 |
| ERK | _ | Rabbit IgG | 9102 | AB_330744 | 1:1000 |
| p-AKT S473 | 193H12 | Rabbit IgG | 4058 | AB_331168 | 1:1000 |
| AKT | _ | Rabbit IgG | 9272 | AB_329827 | 1:1000 |
| p-S6 S240/244 | _ | Rabbit IgG | 2215 | AB_331682 | 1:1000 |
| S6 | 5G10 | Rabbit IgG | 2217 | AB_331355 | 1:1000 |
| BIM | C34C5 | Rabbit IgG | 2933 | AB_1030947 | 1:1000 |
| NOXA | D8L7U | Rabbit IgG | 14766 | AB_2798602 | 1:1000 |
| p-4EBP1 T37/46 | 236B4 | Rabbit IgG | 2855 | AB_560835 | 1:1000 |
| 4EBP1 | 53H11 | Rabbit IgG | 9644 | AB_2097841 | 1:1000 |
| BCL-xL | _ | Rabbit IgG | 2762 | AB_10694844 | 1:1000 |
| BCL-2 | D55G8 | Rabbit IgG | 4223 | AB_1903909 | 1:1000 |
| MCL-1 | D35A5 | Rabbit IgG | 5453 | AB_10694494 | 1:1000 |
| BAX | _ | Rabbit IgG | 2772 | AB_10695870 | 1:1000 |
| BAK | D4E4 | Rabbit IgG | 12105 | AB_2716685 | 1:1000 |
| Cleaved PARP | D64E10 | Rabbit IgG | 5625 | AB_10699459 | 1:1000 |
| Cleaved caspase-3 | _ | Rabbit IgG | 9661 | AB_10699459 | 1:1000 |
| β-tubulin | _ | Rabbit IgG | 2146 | AB_2210545 | 1:3000 |
| Secondary antibody | _ | Anti-rabbit | 7074 | AB_2099233 | 1:3000 |

**Footnotes:** All the antibodies were purchased from Cell Signaling Technology, Danvers, MA.
